# Panpulmonate habitat transitions: tracing the evolution of Acochlidia (Heterobranchia, Gastropoda)

**DOI:** 10.1101/010322

**Authors:** Katharina M. Jörger, Bastian Brenzinger, Timea P. Neusser, Alexander V. Martynov, Nerida G. Wilson, Michael Schrödl

## Abstract

The evolution and diversification of euthyneuran slugs and snails was likely strongly influenced by habitat transitions from marine to terrestrial and limnic systems. Well-supported euthyneuran phylogenies with detailed morphological data can provide information on the historical, biological and ecological background in which these habitat shifts took place allowing for comparison across taxa. Acochlidian slugs are ‘basal pulmonates’ with uncertain relationships to other major panpulmonate clades. They present a unique evolutionary history with representatives in the marine mesopsammon, but also benthic lineages in brackish water, limnic habitats and (semi-)terrestrial environments. We present the first comprehensive molecular phylogeny on Acochlidia, based on a global sampling that covers nearly 85 % of the described species diversity, and additionally, nearly doubles known diversity by undescribed taxa. Our phylogenetic hypotheses are highly congruent with previous morphological analyses and confirm all included recognized families and genera. We further establish an ancestral area chronogram for Acochlidia, document changes in diversification rates in their evolution via the birth-death-shift-model and reconstruct the ancestral states for major ecological traits. Based on our data, Acochlidia originated from a marine, mesopsammic ancestor adapted to tropical waters, in the mid Mesozoic Jurassic. We found that the two major subclades present a remarkably different evolutionary history. The Microhedylacea are morphologically highly-adapted to the marine mesopsammon. They show a circum-tropical distribution with several independent shifts to temperate and temperate cold-habitats, but remained in stunning morphological and ecological stasis since the late Mesozoic. Their evolutionary specialization, which includes a remarkable and potentially irreversible ‘meiofaunal syndrome’, guaranteed long-term survival and locally high species densities but also resembles a dead-end road to morphological and ecological diversity. In contrast, the Hedylopsacea are characterized by morphological flexibility and ecologically by independent habitat shifts out of the marine mesopsammon, conquering (semi-)terrestrial and limnic habitats. Originating from interstitial ancestors with moderate adaptations to the mesopsammic world, they reestablished a benthic lifestyle and secondary ‘gigantism’ in body size. The major radiations and habitat shifts in hedylopsacean families occured in the central Indo-West Pacific in the Paleogene. In the Western Atlantic only one enigmatic representative is known probably presenting a relict of a former pan-Tethys distribution of the clade. This study on acochlidian phylogeny and biogeography adds another facet of the yet complex panpulmonate evolution and shows the various parallel pathways in which these snails and slugs invaded non-marine habitats. Given the complex evolutionary history of Acochlidia, which represent only a small group of Panpulmonata, this study highlights the need to generate comprehensively-sampled species-level phylogenies to understand euthyneuran evolution.

## 1 Introduction

Habitat transition from sea to land or freshwater and vice versa are considered infrequent across all metazoan clades, except for tetrapod vertebrates (Vermeij and Dudley, 2000). The general rarity of habitat shifts in the evolution of the animal kingdom is likely due to the adaptive costs that a new physical environment demands, for example concerning respiration, osmoregulation, reproduction and defensive features, and the competitive disadvantage of the new invaders against well-adapted incumbents (Vermeij and Dudley, 2000; Vermeij and Wesselingh, 2002). The evolution of euthyneuran gastropods, however, defies such generalizations: euthyneuran snails and slugs inhabit all aquatic and terrestrial habitats and pulmonate taxa especially have shown numerous independent shifts between habitats (Barker, 2001; Dayrat et al., 2011; Klussmann-Kolb et al., 2008; Mordan and Wade, 2008). The invasion of new habitats (especially terrestrial ones) played a key role in the diversification of euthyneuran gastropods leading to highest species richness and ecological diversity among this class of Mollusca (Mordan and Wade, 2008). Studying the historical context and the ecological and biological precursors, which facilitated habitat shifts in euthyneuran gastropods, should allow for comparative evaluation to other taxa in the future. Why can barriers preventing habitat transition be overcome more easily by some groups of invertebrates than by others?

To study the evolution of features such as habitat transitions, a robust phylogenetic hypothesis on the sister group relationships among major clades and between taxa inhabiting different environments is required. Unfortunately, until recently, the phylogeny of Euthyneura could not be resolved satisfactorily (Brenzinger et al., 2013; Wägele et al., 2014) via cladistic analyses of morphological markers, probably due to a high degree of homoplasy (Dayrat and Tillier, 2002). Molecular phylogenetics has contradicted the traditional division of Euthyneura in ‘Opisthobranchia’ and ‘Pulmonata’ (Grande et al., 2004; Klussmann-Kolb et al., 2008) and retrieved some former opisthobranch and ‘lower heterobranch’ clades in pulmonate relationships (Jörger et al., 2010b; Klussmann-Kolb et al., 2008; Schrödl et al., 2011a). For continuity in terminology, these newly allied groups are recognized with traditional pulmonates using the taxon name Panpulmonata (Jörger et al., 2010b). The new classification of Euthyneura is consistently retrieved based on molecular ‘standard markers’ (nuclear 18S and 28S rRNA and mitochondrial 16S rRNA and cytochrome oxidase subunit I) (e.g., Dayrat et al., 2011; Dinapoli and Klussmann-Kolb, 2010; Dinapoli et al., 2011; Jörger et al., 2010b; Klussmann-Kolb et al., 2008), but none of these molecular studies provides well-supported sister group relationships among the major panpulmonate taxa. The new classification was not supported by combining data from mitochondrial genomes analyzed for ‘Opisthobranchia’ (Medina et al., 2011) and ‘Pulmonata’ (White et al., 2011). However, mitogenomics does not seem suitable to resolve basal euthyneuran relationships (Bernt et al., 2013; Stöger and Schrödl, 2013). The first phylogenomic approaches support the new classification into Euopisthobranchia and Panpulmonata but also still lack support for inner panpulmonate relationships (Kocot et al., 2013; Zapata et al., 2014). Although considerable advances have been achieved in regards off well-supported monophyly of many major euthyneuran clades, ambiguities in the phylogenetic relationships among the major panpulmonate taxa still hinder overall evolutionary approaches, e.g. to address the transitions between aquatic and terrestrial habitats (Dayrat et al., 2011). Thus, at the current stage of knowledge, the evolution of panpulmonates can only be traced via focusing on well-supported, undisputed clades, adding step-by-step to the complex picture.

The Acochlidia form one of the suitable panpulmonate clades to study habitat transitions. This slug lineage mainly inhabits the interstitial spaces of the marine intertidal and shallow subtidal sands. Based on the current phylogenetic hypothesis on Acochlidia (Jörger et al., 2010b; Schrödl and Neusser, 2010), their evolution comprises several habitat shifts 1) into the mesopsammon (i.e., the habitat of the interstitial spaces of sands *sensu* Remane (1940)) in the putative marine, benthic ancestor and 2) out of the mesopsammon - uniquely for interstitial gastropods - reestablishing a benthic lifestyle in a brackish environment in Pseudunelidae (Neusser and Schrödl, 2009), limnic habitats in Acochlidiidae (see e.g., Brenzinger et al., 2011b; Bücking, 1933; Haynes and Kenchington, 1991; Wawra, 1974, 1979) and Tantulidae (Neusser and Schrödl, 2007; Rankin, 1979) and (semi-)terrestrial habitats in Aitengidae (Neusser et al., 2011a; Swennen and Buatip, 2009). In the past years, detailed microanatomical redescriptions from representatives of all seven acochlidian families and most of the 13 genera were conducted (Brenzinger et al., 2011a; Brenzinger et al., 2011b; Eder et al., 2011; Jörger et al., 2010a; Jörger et al., 2008; Jörger et al., 2007; Kohnert et al., 2011; Neusser et al., 2011a; Neusser et al., 2006; Neusser et al., 2009a; Neusser et al., 2011b; Neusser et al., 2009b; Neusser and Schrödl, 2007, 2009). In combination with a cladistic approach towards the phylogeny of Acochlidia based on a comprehensive matrix of morphological and ecological characters (Schrödl and Neusser, 2010), these studies provide a reliable morphological dataset to study morphological adaptations preceding or resulting from habitat shifts and compare different evolutionary strategies across Acochlidia. Integrative approaches also demonstrated the limits of morphology, however, revealing a high degree of pseudocryptic or fully cryptic speciation in mesopsammic lineages (Jörger et al., 2012; Neusser et al., 2011b). Moreover, convergent adaptations to the mesopsammic habitat were critically discussed as potentially misleading signal in morpho-anatomical phylogenetic analyses (Schrödl and Neusser, 2010).

In the present study, we aim to establish a molecular phylogeny of Acochlidia with a comprehensive panpulmonate outgroup sampling and including all valid acochlidian species. Over the past years and with the support of a series of international collaborators (please see Acknowledgements for details) we successfully recollected at type localities worldwide (see Fig. 1) and are currently able to cover approx. 85 % of the described acochlidian diversity and adding another 50 % previously unknown lineages. In addition to generating this phylogeny, we also aimed to combine carefully calibrated molecular clock analyses and model-based ancestral area reconstruction analyses to retrieve ancestral area chronograms as basis for hypotheses on vicariance events and dispersal in Acochlidia. Moreover, we wanted to reconstruct the history of major ecological traits (habitat, lifestyle and climate) by inferring ancestral states. Based on phylogenetic trees the rate of evolution that created present-day diversity can be evaluated. Typically, rates of diversification changes (speciation minus extinction) are detected via slope changes in lineage-through-time plots (LTT). We also wanted to implement a recent powerful likelihood approach that can detect shifts in the rate of diversification and avoid stochastic errors e.g. when only a small number of specimens is available (Stadler, 2011a, see material and methods for details). Moreover, it can account for incomplete taxon sampling (Stadler, 2011a), a fact which needs to be considered in acochlidian evolution with the vast majority of the world’s mesopsammic habitat still unsampled (Jörger et al., 2012; Neusser et al., 2011b).

**Figure 1:**
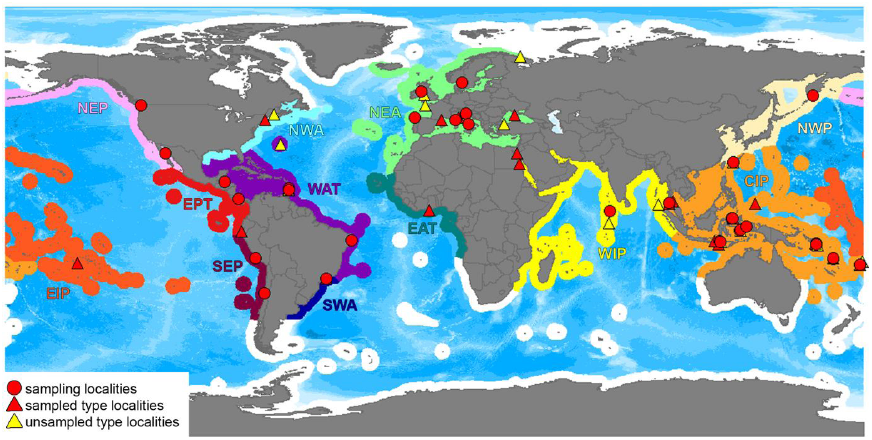
Sampling localities of Acochlidia. Triangles indicate type localities (when colored in red material from type locality was included in the study, yellow triangles: unsampled type localities), red dots represent collecting sites of material included in this study (for collectors, coordinates and details on the collecting sites, see Additional material 1). Yellow triangles: unsampled type localities. Biogeographic areas modified after system of Marine Ecoregions of the world (MEOW) by Spalding et al (2007), using their original shape-file to generate the map in DIVA-GIS. CIP – Central Indo-Pacific; EAT – tropical Eastern Atlantic; EIP – Eastern Indo-West Pacific; EPT – tropical Eastern Pacific; NEA – North-Eastern Atlantic (+Mediterranean+Black Sea); NEP – North-Eastern Pacific; NWA – North-Western Atlantic; NWP – North Western Pacific; SEP – South-East Pacific; SWA – South-Western Atlantic; WAT – tropical Western Atlantic; WIP – Western Indo-Pacific. White coastlines mark unsampled biogeographic regions.

With this combination of phylogenetic approaches to reconstruct the evolutionary history of Acochlidia, we aim to provide knowledge on the biogeographic background of habitat shifts in Acochlidia and reevaluate morphological, biological and behavioral traits which might have triggered or resulted from these transitions. The role of progenesis in the evolution of this meiofaunal taxon is discussed, as is the impact of secondary ‘gigantism’ to cope with the physical requirements in freshwater and terrestrial habitats. The conclusions on the evolutionary history of Acochlidia aim to add a piece to the complex puzzle of the evolution and radiation of hyperdiverse panpulmonates and provide insights on requirements and consequences of habitat shifts in marine invertebrates.

## 2 Material and methods

### 2.1 Sampling and fixation

Sampling effort for Acochlidia was conducted in intertidal and subtidal sands worldwide, whenever possible covering type localities of nominal species (see Fig. 1 and Additional material 1 for collecting sites). In total, 36 described and 30 undescribed lineages (identified as molecular operational taxonomic units –MOTUs) are included here. Limnic specimens were collected by hand from the undersides of stones in rivers and streams. All material was observed alive and whenever possible photographed in the field. Limnic specimens were anesthetized prior to fixation using menthol crystals. Meiofaunal specimens were extracted from sand sampled using a MgCl_2_-seawater solution and careful decantation technique (Jörger et al., 2014; Schrödl, 2006) and again anesthetized with MgCl_2_ prior to fixation to prevent retraction of the head-foot complex. Material was fixed in 75 % ethanol (for preparation of hard structures, such as radulae, spicules and copulatory stylets), 96-99 % ethanol (molecular analyses) and 2.5-4 % glutardialdehyde buffered in cacodylate (for histology and ultrastructure).

Unfortunately, our analyses lack one family of Acochlidia: monotypic Tantulidae. *Tantulum elegans* Rankin 1979 was discovered by Rankin (1979) in the muddy interstices of a freshwater mountain spring of the Caribbean Island of St. Vincent. The original type material is unsuitably fixed for sequencing attempts and was not obtained from the Royal Ontario Museum. New recollection attempts at the type locality (by KMJ) in February 2009 failed. The described locality could be localized precisely based on the available literature (Harrison and Rankin, 1976; Rankin, 1979), but has changed considerably probably due to agriculture and the (only) spring where *Tantulum* occurred had disappeared at least during that time of year.

### 2.2 DNA extraction, amplification and sequencing

DNA was extracted from entire specimens (in minute meiofaunal taxa) or from foot tissue (limnic taxa) using the DNA Qiagen Blood and Tissue Kit or the Macherey-Nagel NucleoSpin Tissue Kit. We followed the manufacture’s extraction protocol, with overnight tissue lysis. We amplified four genetic standard markers via polymerase chain reaction (PCR): mitochondrial cytochrome c oxidase subunit I (COI) and 16S rRNA, and nuclear 18S rRNA and 28S rRNA using the same protocols and primers as listed in Jörger et al. (2010b). Successful PCR products were cleaned up using ExoSap IT (Affymetrix Inc.) (COI, 16S rRNA) or Macherey-Nagel NucleoSpin Gel and PCR Clean Up (28S rRNA). Cycle sequencing and sequencing reactions were performed by the Sequencing Service of the LMU using the PCR primers, Big Dye 3.1 and an ABI capillary sequencer. Voucher specimens are deposited at the Bavarian State Collection of Zoology (ZSM, Munich), DNA aliquots are stored and publicly available through the DNAbank network (http://www.dnabank-network.org) and all sequences are deposited to GenBank at NCBI (http://www.ncbi.nlm.nih.gov/genbank/) (see Table 1 for accession numbers).

**Table 1:**
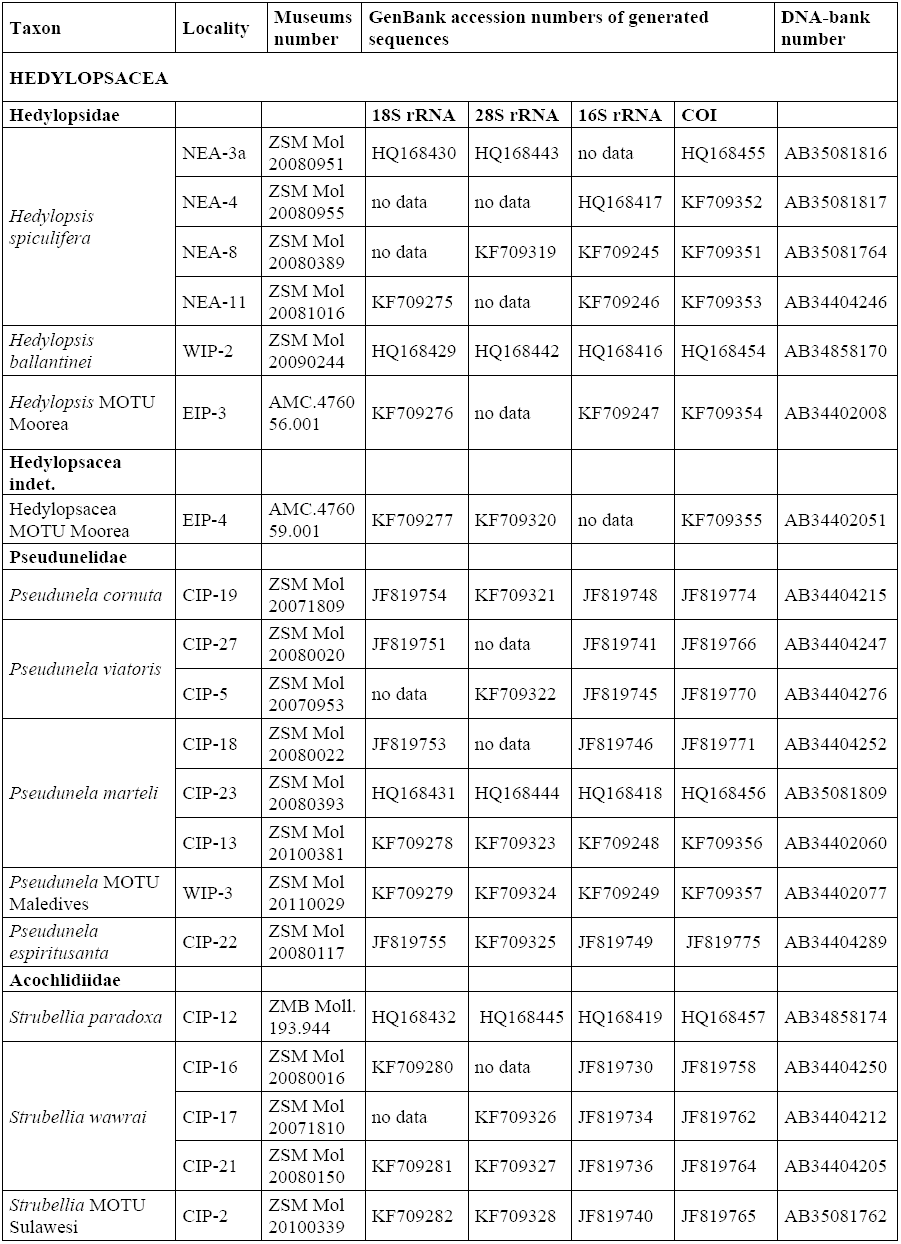
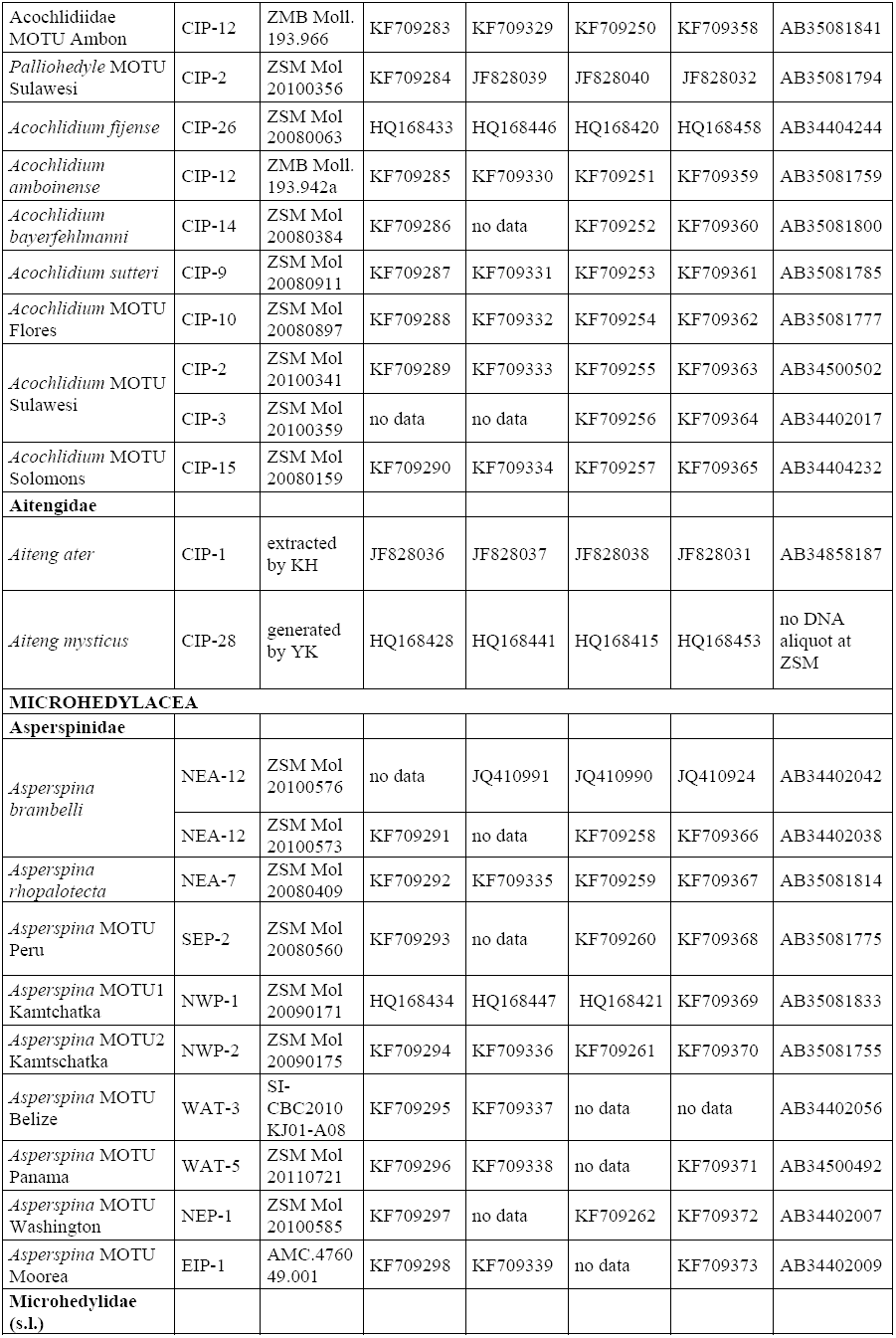
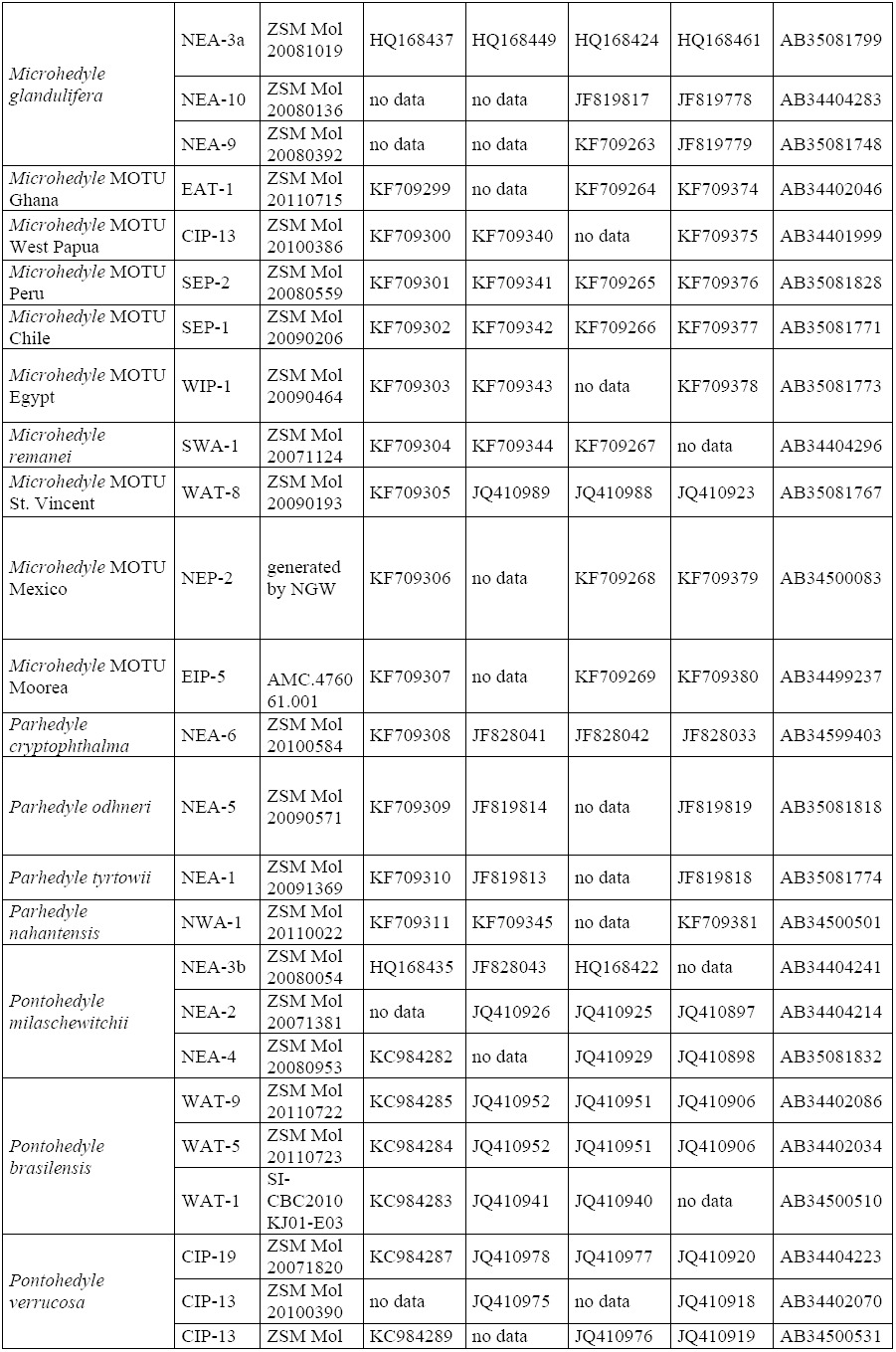
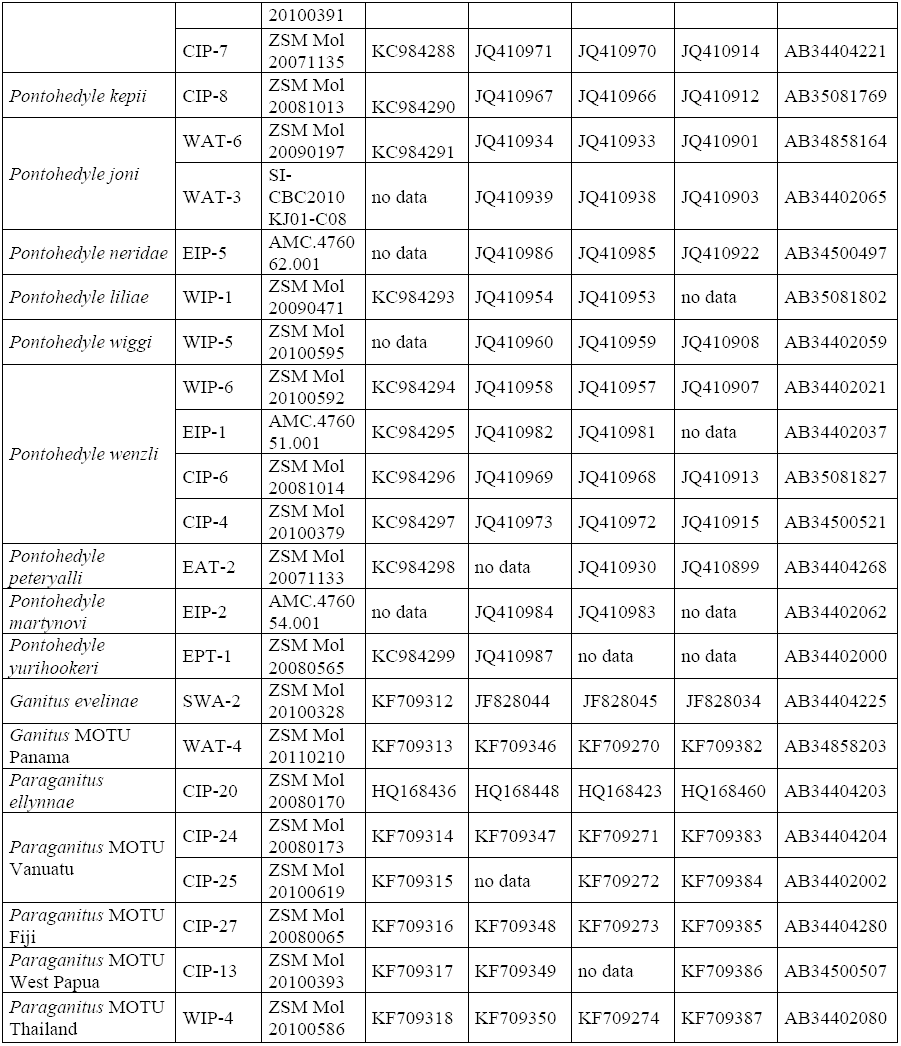
List of museums numbers of voucher material, DNA voucher numbers (all ZSM) and GenBank numbers of the material of Acochlidia analyzed in the present study. * marks sequences generated within this study, all remaining sequences were retrieved from GenBank. AM – Australian Museum, Sydney; SI – Smithsonian Institute, Washington, D.C.; ZSM – Bavarian State Collection of Zoology, Munich. DNA and sequences generated by KMJ if not stated otherwise: KH – Katharina Händeler – University of Bonn; YK – Yasunori Kano, University of Tokyo; NW – Nerida Wilson, Scripps Institution of Oceanography, La Jolla.

### 2.3 Phylogenetic analyses

Sequences were edited with BioEdit (Hall, 1999) and Geneious 5.2 (Drummond et al., 2010). All sequences generated in this study were checked for putative contamination using BLAST searches (Altschul et al., 1990) to compare our sequences with published sequences in GenBank (http://blast.ncbi.nlm.nih.gov). Outgroup selection followed the latest phylogenetic hypothesis on the origin of Acochlidia (Jörger et al., 2010b), aimed to cover all major lineages of Panpulmonata and also included more distantly related Euopisthobranchia and a ‘lower heterobranchs’. Outgroup sequences were retrieved from GenBank (see Table 2 for accession numbers). Alignments for each marker were generated using MAFFT (E-ins-I-option) (Katoh et al., 2005). Ambiguous positions in the alignment were removed using Aliscore (Misof and Misof, 2009). Comparative masking with Gblocks (Talavera and Castresana, 2007) (options for a less stringent selection) resulted in the removal of 300 more nucleotide positions but the topology of the final ML-tree was not affected. The COI alignment was checked manually and by translation into amino acids for potential shift in the reading frame and for stop codons. Sequences were concatenated using Sequence Matrix (Meier et al., 2006), and BioEdit (Hall, 1999) for the translated COI alignment.

**Table 2:**
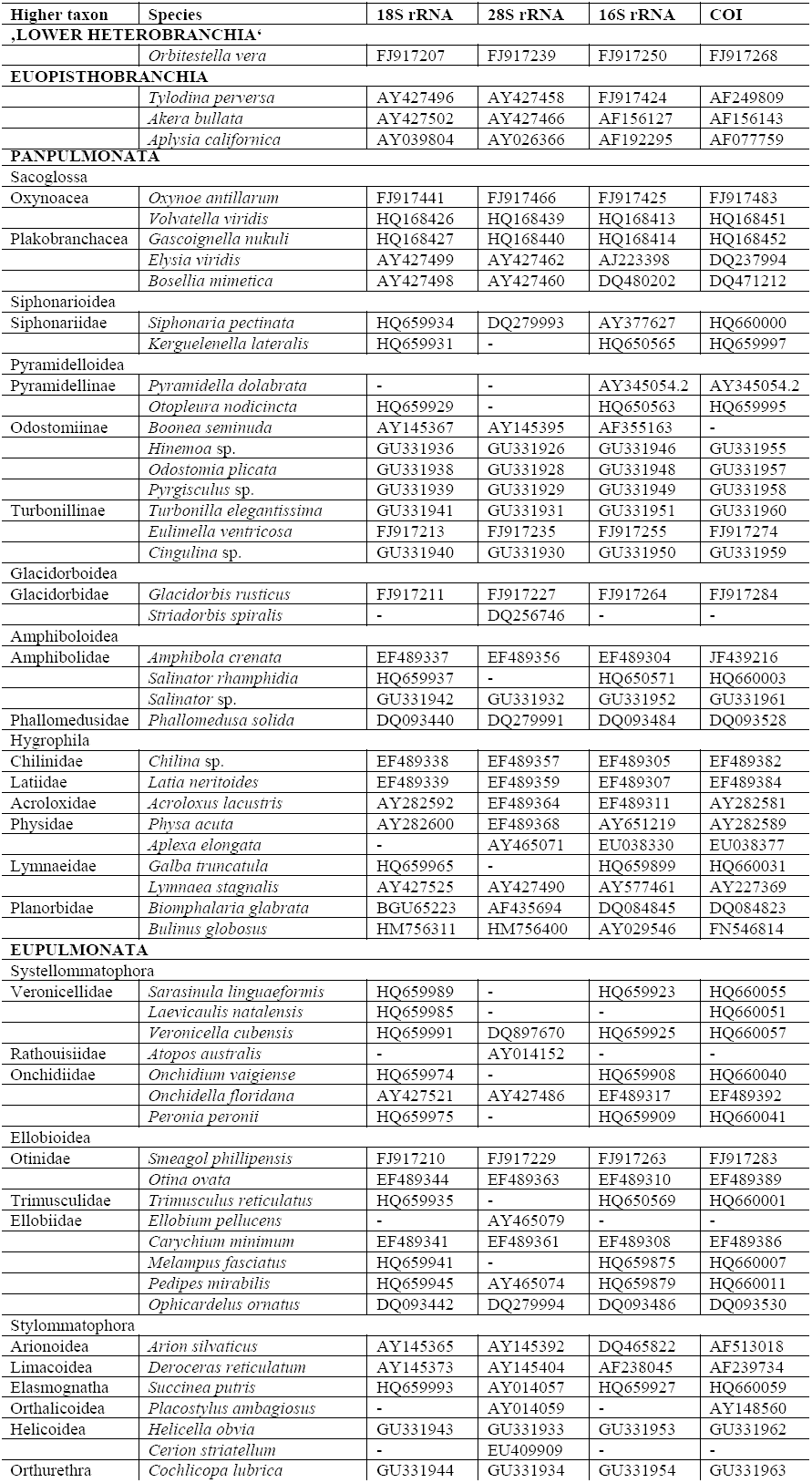
List of included panpulmonate outgroup taxa for phylogenetic analyses with GenBank accession numbers.

Models were selected using jModeltest (Posada, 2008) from 5 substitution schemes and 40 models for each individual marker and the concatenated dataset. This resulted in the GTR+gamma+I-model for all four markers. We conducted maximum-likelihood analyses of the concatenated for-marker dataset, analyzed in four partitions corresponding to each marker. In our RAxMLv7.2.8 analyses we followed ‘the hard and slow way’ in RAxMLv7.0.4 Manual and generated five parsimony starting trees, applied 10 different rate categories, 200 random starting trees and non-parametric bootstrapping on 1000 trees. *Orbitestella vera* (Powell 1940) (Orbitestellidae, ‘lower Heterobranchia’) was defined as the outgroup.

### 2.4 Molecular dating

Molecular clock analyses – To estimate divergence times in Acochlidia we conducted relaxed molecular clock analyses using BEAST v1.6.1 (Drummond and Rambaut, 2007) on our concatenated four marker dataset. All analyses were run with the relaxed uncorrelated lognormal model in four partitions corresponding to each marker and under the Yule process using the GTR+Gamma+I substitution model for each marker. We selected calibration points, which we considered most reliable (i.e. taxa with a decent fossil record and comparably reliable assignation to present day taxa) and most conservative (i.e. potential older representatives with controversial assignment were ignored). A gamma-shaped distribution prior was selected setting a hard prior on the minimum age of the respective node, highest probability was assigned to the ages selected from literature and a 10 % range was chosen for the distribution curve. Fossil timing follows Tracey et al. (1993) and Bandel (1994) using five calibration points: the minimum age of the Anaspidea was set to 190 mya, Ellobiidae to 145 mya, Siphonariidae to 150 mya and Lymnaeidae to 140 mya. Moreover, we calibrated the closure of the Isthmus of Panama to 2.1 mya, accommodating the possibility of subsequent submersion after the closure about 3.4 mya (Cox and Moore, 2010). Analysis with all five calibration points was run for 100 000 000 generations. Results were analyzed in Tracer v1.5, and all values reached good effective sampling sizes. To produce a consensus tree, all trees were combined in TreeAnnotator v1.6.1, with the first 10 000 trees discharged as burn-in. To evaluate the impact of each calibration point on the time estimations, we performed sensitivity analyses and ran five additional BEAST analyses (30 000 000 generations each) omitting one calibration point in each of the analyses.

Rate of evolution – we applied two different methods to detect changes in the rate of evolution of Acochlidia. As input tree we used the ultrametric tree from our molecular clock analyses in BEAST and removed outgroups and multiple individuals of one species prior to analyses in R. We generated lineage-through-time plots (LTT) in R using the ape package (Paradis et al., 2004). Additionally, we conducted maximum-likelihood analyses applying the birth-death-shift process (Stadler, 2011a) to infer diversification rate changes in TreePar (Stadler, 2011b). As described by Stadler (2011a) the likelihoods of the model allowing *m* rate shifts were compared to the model allowing *m+1* rate shifts via the likelihood ratio test, applying a 0.99 criterion.

### 2.5 Ancestral area reconstruction

We used two different algorithms of ancestral area reconstruction: Statistical Dispersal-Vicariance analysis (S-DIVA) in RASP 2.1b (Yu et al., 2012) and dispersal-extinction-cladogenesis model (DEC) implemented in Lagrange (Ree et al., 2005; Ree and Smith, 2008). We coded the areas based on the biogeographic realms of the Marine Ecosystem of the World (MEOW) system designed for the world’s coastal and shelf areas (Spalding et al., 2007). In general, meiofaunal Acochlidia lack a planktonic larval stage, thus dispersal abilities are considered low (Jörger et al., 2012; Neusser et al., 2011b; Swedmark, 1968). Additional to the MEOW realms, we further subdivided Eastern/Western regions of the Atlantic and Pacific, and separating the Pacific and Atlantic Coast of temperate South America, which resulted in 12 coded areas for the included material: Western Indo-West Pacific (WIP), central Indo-West Pacific (CIP), Eastern Indo-West Pacific (EIP), tropical Eastern Pacific (EPT), Southeast Pacific (SEP), Northeast Pacific (NEP), Northwest Pacific (NWP), tropical Western Atlantic (WAT), Northwest Atlantic (NWA), Southwest Atlantic (SWA), tropical Eastern Atlantic (EAT), Northeast Atlantic (NEA) (see Figure 1). No subdivisions into provinces of the MEOW were adopted, as they contradict distribution ranges based on molecular analyses and haplotype networks in comparably wide spread microhedylacean Acochlidia (see Eder et al., 2011; Jörger et al., 2012). Ranges of ancestral areas should resemble those of recent representatives (Clark et al., 2008, Hausdorf, 1998), we thus allowed for a maximum of two areas at each ancestral node. For the input tree in both analyses, we used our chronogram generated with BEAST using all calibration points discussed above, with outgroups removed. Multiple individuals from one species distributed in the same area were removed for ancestral area reconstruction. In Lagrange analyses, we included five dispersal matrices to accommodate differences in dispersal probabilities during the changing geological history of Earth: 1) from mid Jurassic (175 mya) to early Cretaceous (120 mya) prior to the breakup of Gondwana; 2) Early Cretaceous (120 mya) to late Cretaceous (90 mya) as the circum Tethyan Seaway allowed worldwide exchange of tropical coastal faunas, and the South Atlantic formed; 3) late Cretaceous (90 mya) to Miocene (20 mya) characterized by the ongoing Tethyan Seaway and completed formation of the Atlantic and Beringia prohibiting exchange between Northwest Pacific and North Atlantic fauna; 4) Miocene (20 mya) to Pliocene (3.5 mya): closure of Tethyan Seaway isolates Indo-West Pacific from Atlantic/ Pacific fauna, Bering Strait opens; 5) Pliocene (3.5 mya) to present: closure of the Panamanian Isthmus separates Western Atlantic from Eastern Pacific. Dispersal probabilities range from 0.1 between disconnected, separated areas and 1.0 between directly adjacent coastal areas. Dispersal between areas separated by one intermediate realm was set to 0.25, as were adjacent areas with supposed dispersal barriers for Acochlidia (e.g. trans-oceanic ranges or present day East Pacific barrier). For comparison, we ran a more constrained analyses restricting dispersal to (directly) adjacent regions.

### 2.6 Transitions in ecological traits

We reconstructed ancestral states of ecological traits (climate: tropical, temperate, temperate-cold; habitat: marine, amphibious-marine, brackish, limnic; life style, interstitial, benthic) using MESQUITE 2.75 (Maddison and Maddison, 2011) on our fully resolved maximum-likelihood topology. For comparison, we also repeated analyses using the slightly different topology generated in Bayesian analyses conducted with BEAST. To assess ancestral states, we performed maximum-parsimony and maximum-likelihood analyses, in the latter applying the Mk1 model (assuming all changes between states are equally probable). The selection of a more complex model was not supported using the likelihood ratio test from results in Bayes Traits (Pagel and Meade, 2006).

## 3 Results

### 3.1 Phylogenetic analyses

We retrieved our hypothesis on the phylogeny of Acochlidia from a maximum-likelihood analysis conducted with RAxML 7.2.8 on the concatenated four marker dataset (nuclear 18S rRNA, 28rRNA and mitochondrial 16S rRNA and COI) analyzed in four partitions corresponding to each marker. An overview of the topology with the sister group relationships of Acochlidia and the relationships among acochlidian families, is provided in Fig. 2. We included 57 outgroup taxa resembling a broad range of panpulmonate taxa to resolve the origin of Acochlidia within Panpulmonata. Unfortunately, as in previous analyses (Jörger et al., 2010b; Klussmann-Kolb et al., 2008) none of the sister group relationships of major panpulmonate subgroups is statistically supported (i.e. receives bootstrap values (BS) ≥ 75). But in all analyses conducted herein Acochlidia are monophyletic and are sister to a clade comprised of Hygrophila+(Pyramidelloidea+(Glacidorboidea+Amphiboloidea)). We included 36 out of 43 valid acochlidian species and 30 individuals belonging to roughly 20 or more putatively undescribed species, identified here as molecular operational taxonomic units (MOTUs) (see Fig. 3). The included taxa represent all recognized acochlidian families and genera, apart from the monotypic Tantulidae, with *Tantulum elegans* Rankin, 1979 being unavailable for molecular analyses. All acochlidian (super-)families inferred from cladistic analyses of morphological characters (Schrödl and Neusser, 2010) resulted monophyletic. In concordance with morphological analyses, Acochlidia split into Hedylopsacea (comprised of Hedylopsidae+(Aitengidae+(Hedylopsacea *incertae sedis*+(Pseudunelidae +Acochlidiidae)))) and Microhedylacea (comprised of Asperspinidae+Microhedylidae s.l. (i.e. including paraphyletic Ganitidae)). Neither the sistergroup relationship of Hedylopsacea and Microhedylacea, nor the deep splits within these clades, however, are supported by BS ≥ 75. Most recognized acochlidian families receive high statistical support (BS 75-100), but not Pseudunelidae and Acochlidiidae (see Fig. 3). One undescribed species of Hedylopsacea, which also unites a unique combination of morphological characters (KMJ unpublished data), resulted as the sister clade of Pseudunelidae+Acochlidiidae and thus formed a still unnamed ‘family-level’ clade (Hedylopsacea MOTU Moorea in Fig. 3). The topology of our analyses was concordant between the different phylogenetic approaches (maximum-likelihood, Figs. 2,3; Bayesian inference, Fig. 4) with the exception of amphibious Aitengidae, either being sister to all remaining hedylopsacean taxa (Fig. 4) or sister to Pseudunelidae+(Hedylopsacea sp.+Acochlidiidae) (Figs. 2,3). In our analyses all recognized acochlidian genera were monophyletic, apart from *Microhedyle*, which is paraphyletic due to the inclusion of *Paraganitus*, a result concordant with morphological analyses, but in the latter it is paraphyletic due to inclusion of Ganitidae (*Paraganitus*+*Ganitus*) (Schrödl and Neusser, 2010). With the exception of the genus *Pseudunela*, all monophyletic acochlidian genera are statistically highly supported (see Fig. 3). Next to the undetermined clade ‘Hedylopsacea sp.’, another undescribed species ‘Acochlidiidae sp.’, forms a separate ‘genus-level’ clade sister to *Palliohedyle*+*Acochlidium*. All remaining MOTUs cluster within recognized genera.

**Figure. 2:**
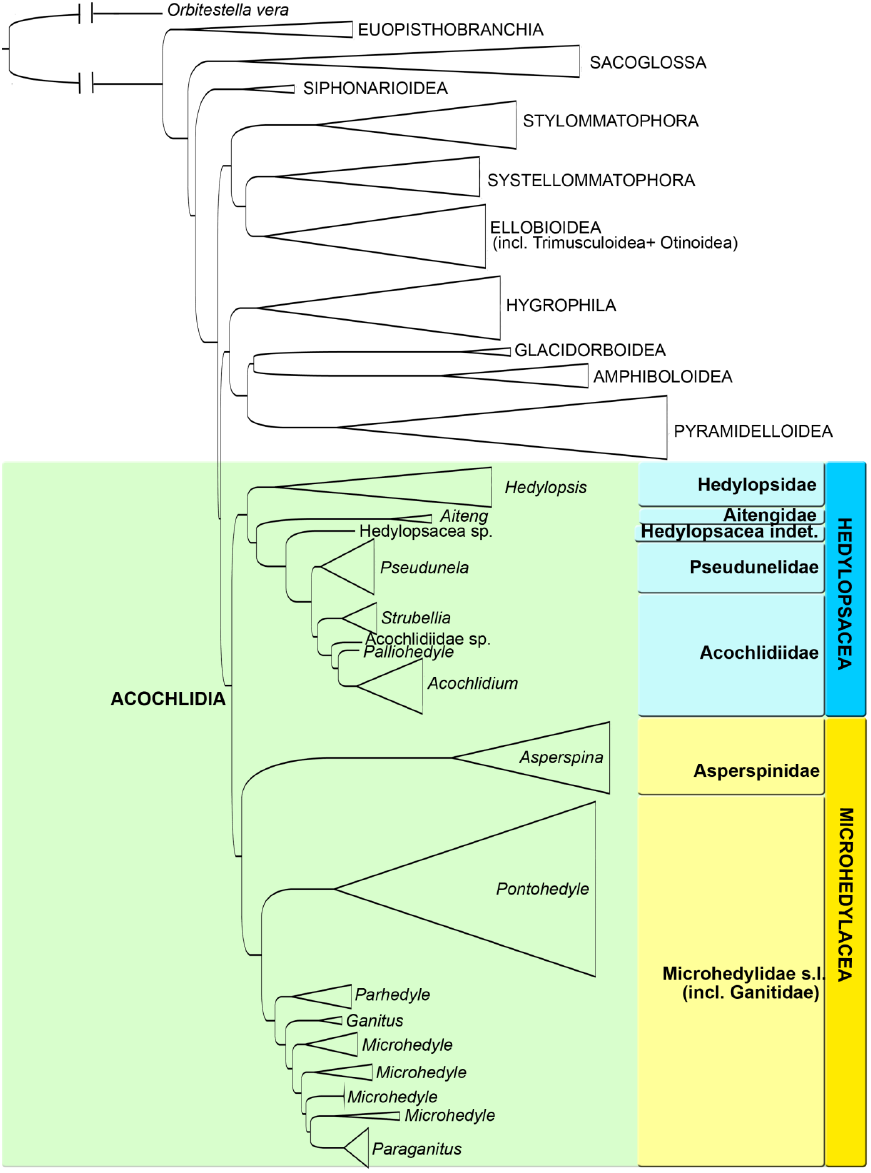
Overview of the phylogenetic relationships of Acochlidia within Panpulmonata, based on maximum likelihood analyses of the concatenated four marker dataset (mitochondrial COI and 16S rRNA and nuclear 28S rRNA and 18S rRNA).

**Figure. 3:**
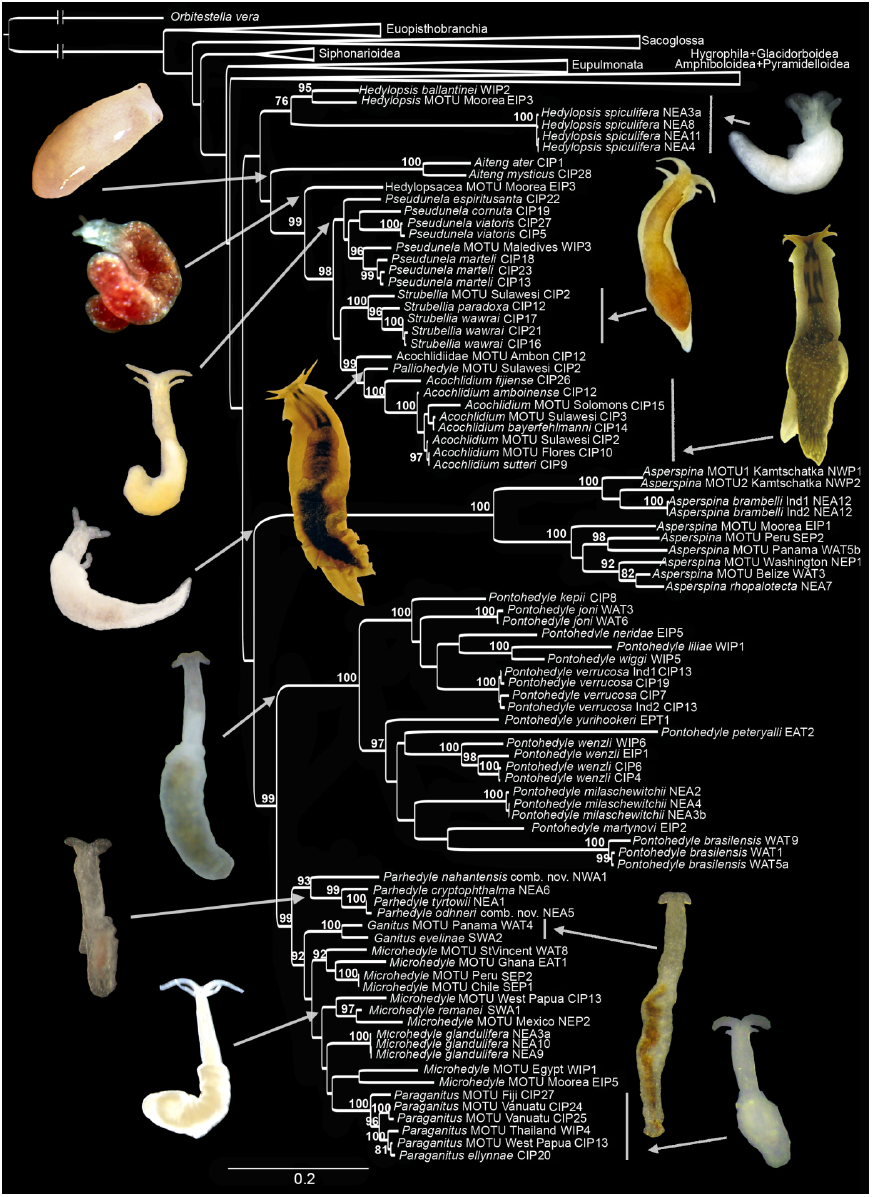
Phylogeny of Acochlidia shown to species level (outgroups collapsed for presentation purposes), based on maximum-likelihood analyses of the concatenated four marker dataset (mitochondrial COI and 16S rRNA and nuclear 28S rRNA and 18S rRNA).

### 3.2 Molecular dating and rate of evolution

We performed sensitivity analyses by six independent molecular clock analyses to test the effect of different calibration points to our dataset (see material and methods for details). Differences for our time estimates on acochlidian evolution only vary slightly among the different analyses and no general pattern could be observed that one calibration point rejuvenates or artificially ages the recovered time estimates (see Additional material 2 for resulting node ages and ranges of 95 % highest posterior densities (HPD)). The values reported below and presented in Fig. 4 refer to our main molecular clock analyses using all calibration points listed in the material and method section. For the entire chronogram with uncollapsed outgroups see Additional file 3. According to our data, most major panpulmonate clades originated in the Mesozoic Jurassic (i.e. Siphonarioidea, Sacoglossa, Stylommatophora, (Systellommatophora+Ellobioidea), Hygrophila, (Glacidorboidea+Amphiboloidea), Pyramidelloidea and Acochlidia), the split between Systellommatophora/Ellobioidea and the split between Glacidorboidea/Amphiboloidea are dated to the early Cretaceous (see Additional file 3). The origin of Acochlidia is dated to the mid Jurassic 176.6 mya (HPD: 207.9–171.0 mya). The diversification into the two major acochlidian clades – Hedylopsacea and Microhedylacea – occurred shortly after but still in Jurassic times 169.2 mya (HPD: 200.9–159.9 mya). The diversification of the Hedylopsacea into the recent families started in the late Jurassic to early Cretaceous with the origin of Aitengidae estimated to 144.4 mya (175.4–97.0 mya), the origin of Hedylopsidae to 121.8 mya (HPD: 160.1–83.5 mya) and the split between Hedylopsacea *incertae sedis* and (Pseudunelidae+Acochlidiidae) estimated to 88.9 mya (HPD: 113.0–48.5mya). Pseudunelidae and Acochlidiidae originate in the Paleogene 33.7 mya (HPD: 58.3–32.0 mya). Genera of Acochlidiidae originate during late Paleogene/early Neogene: the origin of *Strubellia* is estimated to 29.6 mya (HPD: 50.6–27.1 mya), the origin of undescribed Acochlidiidae sp. To 24.3 mya (HPD: 40.5–19.9 mya), and the origin of *Palliohedyle* and *Acochlidium* to 20.6 mya (HPD: 34.4–16.2 mya). The valid microhedylacean families Asperspinidae and Microhedylidae are estimated to have a Mesozoic origin at 165.8 mya (HPD: 193.3–150.4 mya), with the radiation of *Asperspina* and *Pontohedyle* starting in the Cretaceous 84.5 mya/104.3 mya (HPD: 113.7–71.3 mya/133.5–83.5 mya). The origin of *Parhedyle* is estimated to the late Cretaceous/early Paleogene (56.3 mya; HPD: 87.5–44.6 mya). The origin of *Ganitus* and *Paraganitus* is estimated to the late Paleogene 43.2 mya resp. 26.6 mya (HPD: no estimate for *Ganitus* resp. 41.7–20.2 mya for *Paraganitus*).

**Figure. 4:**
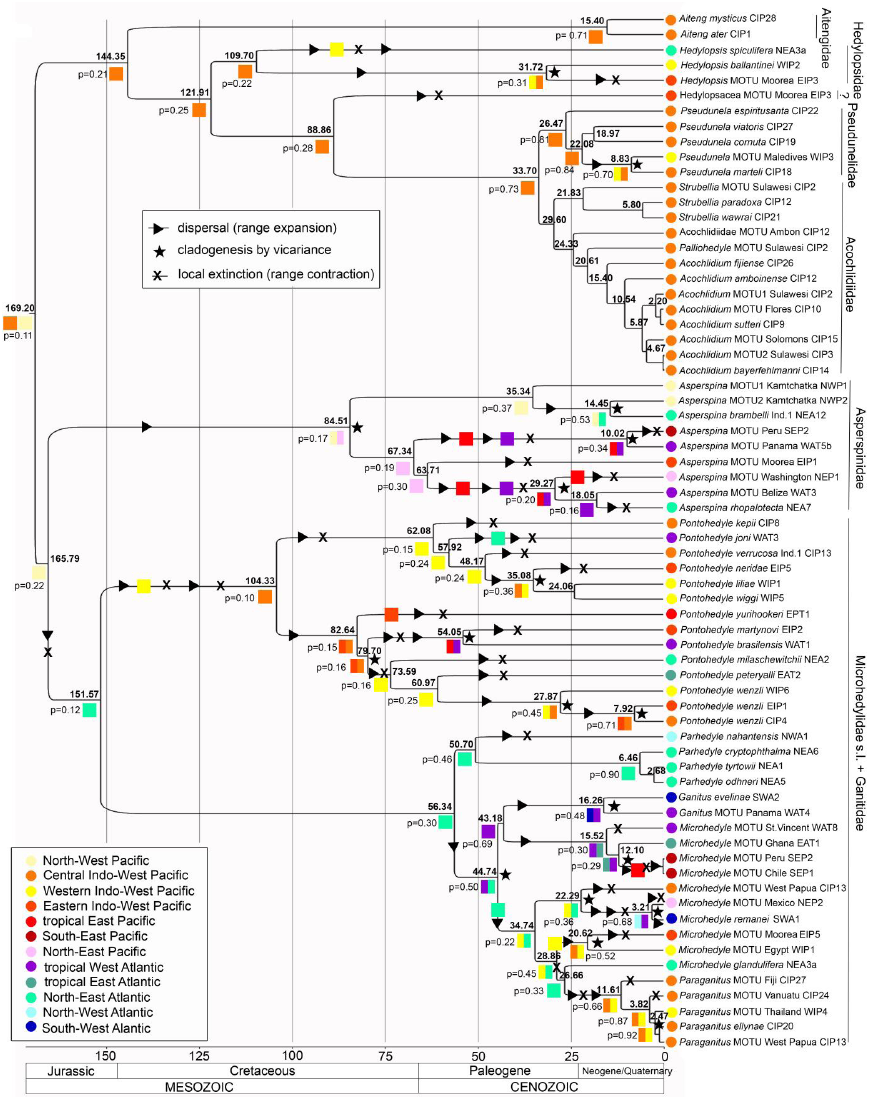
Ancestral area chronogram of Acochlidia, outgroups collapsed for presentation purposes (for complete chronogram see Additional material 3). Divergence times obtained from BEAST v1.6.1 under a relaxed uncorrelated clock model, node ages indicated above nodes. Geological timescale is based on the International Stratigraphic Chart by the International Commission on Stratigraphy (2012). Colored dots at terminals indicate geographic areas occupied the sampled specimens, squares at nodes ancestral area which received highest relative probability (p) in DEC-analyses.

To detect changes in the rate of diversification of Acochlidia we reconstructed a lineage-through-time (LTT) plot. Ignoring the time period prior to 100 mya (because the log number of lineages is too low), the LTT showed a continuous slope with no major changes (see Fig. 5A). One minor period of stasis was present at 40–35 mya followed by a slight increase in diversification, abating again slightly at 10 mya till present. Additionally, we used the likelihood approach with the birth-death-shift process described by Stadler (2011a) on our phylogeny to determine rate changes in the diversification of Acochlidia through time. In all tested scenarios, diversification rates were low with minimum 0.02 and maximum 0.12 per million years. Testing sampling intensities between 90-25 % the likelihood ratio test supported no shifts in rate changes (see Fig. 5B). Diversification rates remained constant between 0.021–0.025 per million years. With sampling intensities lowered to 10 (and 5 %), the model supported one rate shift in the evolution of Acochlidia at approximately 37 mya with the rates of diversification increasing from 0.026 (0.030) to 0.061 (0.075). An alternative scenario testing 1 % sampling intensity also supported one rate shift in evolution of Acochlidia but resulted in a considerable decrease (from 0.118 to 0.024) at approximately 123 mya (see Fig. 5B).

**Figure. 5:**
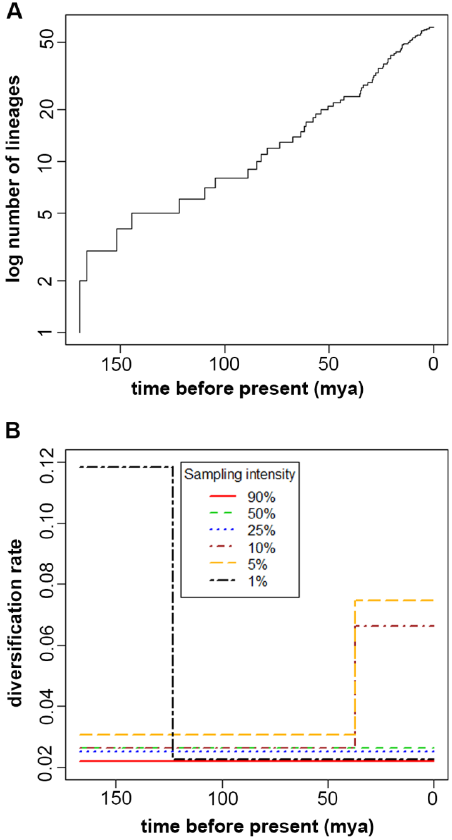
Analyses of shifts in diversification rate of Acochlidia. a) Lineage through time plot, b) birth-death model calculated with TreePar.

### 3.3 Ancestral area reconstruction of Acochlidia

Independent of the chosen approach (DEC model in Lagrange or S-DIVA), or the different models of dispersal allowed between the defined ocean ranges, no ancestral areas with robust support values could be recovered for the basal nodes in Acochlidia. In all approaches, their evolutionary history involves numerous dispersal (range expansion), vicariance and local extinction (range contraction) events. In Lagrange, the less constricted model (favoring dispersal via neighboring areas but not entirely prohibiting dispersal between unconnected areas, see Material and Methods) received better likelihood values and is presented in Fig. 4. Since the S-DIVA analysis was conducted without restrictions on area distributions, several ancestral areas include ‘impossible’ distribution ranges (across unconnected areas). Thus, S-DIVA results are only reported when not contradicted by geography. The paleotopology of the continents changes considerably throughout the evolutionary history of Acochlidia and especially modern Ocean ranges are difficult to assign to Mesozoic ranges. Recent Acochlidia are restricted to costal and shelf areas, we thus aim to approximately allocate recent continental margins to the historic topologies at the ages of the reconstructed nodes (see also Discussion). The North-West Pacific (NWP) and the central Indo-West Pacific (CIP) both receive comparable support values in DEC analyses (relative probability p_r_=0.11 each, referring to the North-Eastern part of the Tethys Ocean and the Western margin of the Panthalassic/ Pacific Ocean) as ancestral area for the diversification of Acochlidia (S-DIVA additionally includes the Eastern Indo-Pacific (EIP) and Western Atlantic Ocean (WAT) as potential ancestral areas). The radiation of Hedylopsacea originated in the waters nowadays belonging to the CIP (Eastern Tethys Ocean - p_r_=0.21 S-DIVA: 0.5 for CIP, 0.5 for CIP+EIP). One major dispersal event occurred at the base of Hedylopsidae via the Western Indo-Pacific (WIP, i.e. Western part of the Tethys) into the North Atlantic (NEA) and also to EIP, in the Cretaceous all well-connected via the circum-equatorial Tethyan Seaway. Based on DEC, two vicariance events (in *Hedylopsis* and *Pseudunela*) occurred isolating sister species in WIP and CIP.

The biogeographic history of Microhedylacea is highly complex and characterised by numerous dispersal, local extinction and vicariance events (see Fig. 4). The primary radiation of Microhedylacea most likely (p_r_=0.22) occurred in the waters nowadays belonging to the NWP (Western part of the Panthalassas/ Pacific Ocean, S-DIVA: 0.25 for CIP or WAT) with an eastward dispersal and range expansion to the North-East Pacific in the stemline of Asperspinidae. A vicariance event split the two asperspinid clades with the NWP-clade dispersing westward around the Asian continent into the North Atlantic. The NEP asperspinid clade (p_r_=0.37, S-DIVA: 1.0 NEP) dispersing south-eastward via the East Pacific (EPT) into the Western Atlantic (WAT) and back again in *Asperspina* MOTU Washington (NEP) and from WAT eastward to NEA. The basal radiation of Microhedylidae s.l. most likely occurred in NEA (in late Jurassic resembled by the forming Atlantic Ocean, p=0.12). Along the stemline of *Pontohedyle* two major dispersal events occurred eastward via WIP into CIP (i.e. through the western part of the Tethys to the eastern part). One of the two major *Pontohedyle* clades disperses via the WIP and from this western part of the Tethys Ocean westward via the North East Atlantic into the Western Atlantic and also eastward into the Eastern Indo-West Pacific (see Fig. 4). The other shows a similar complex picture with several dispersals events among the waters of the Tethyan Seaway in both directions westward and eastward. Our data for example indicates three independent dispersal events in *Pontohedyle* into EIP: twice eastward via CIP and once westward from the East Pacific. The genus *Parhedyle* (restricted to the North Atlantic) most likely (p_r_=0.30) originated in NEA and dispersed into NWA (ancestral area reconstructed with S-DIVA: 1.0 NEA+NWA). A vicariance event split the ancestral population of (*Ganitus*+‘*Microhedyle*’) and (‘*Microhedyle’*+*Paraganitus*), which was distributed across the Atlantic (p_r_=0.50) into an originally Western Atlantic clade including *Ganitus* (p_r_=0.50, S-DIVA: 1.0 WAT) (with one dispersal event back to the tropical East Atlantic and one into the Pacific) and one clade which originated in NEA+WIP (p_r_=0.22) and dispersed westward via WAT to SWA and into the North-East Pacific and three times independently to the East (e.g. along the stemline of *Paraganitus,* with WIP+CIP as ancestral area of its radiation p_r_=0.50, S-DIVA: 1.0 CIP).

### 3.4 Ancestral state reconstruction of ecological traits

We performed ancestral state reconstructions of three major ecological traits (climate, habitat and life style) using maximum-likelihood (values derived from maximum-parsimony are reported when differing). We also coded the traits in our outgroup taxa, but results might not be representative for these clades due to limited taxon sampling (see discussion). Based on our analyses the Acochlidia derive from a marine (likelihood (lh) 0.96) and benthic (lh 0.99) common ancestor with Hygrophila+(Pyramidelloidea+(Glacidorboidea+Amphiboloidea)) and originated in temperate waters (lh 0.98) (see Fig. 6). Along the acochlidian stemline, the ancestral acochlid inhabited tropical waters (lh 0.97) and invaded the interstitial habitat (lh 0.83; most parsimonious state: benthic or interstitial). Rerunning the ancestral state reconstruction without outgroup taxa, the ancestor of Acochlidia was clearly (lh 0.99) marine, interstitial, and inhabiting tropical waters. This also affects likelihoods for the hedylopsacean radiation originating from a marine, mesopsammic and tropical ancestor (lh 0.99).

**Figure. 6:**
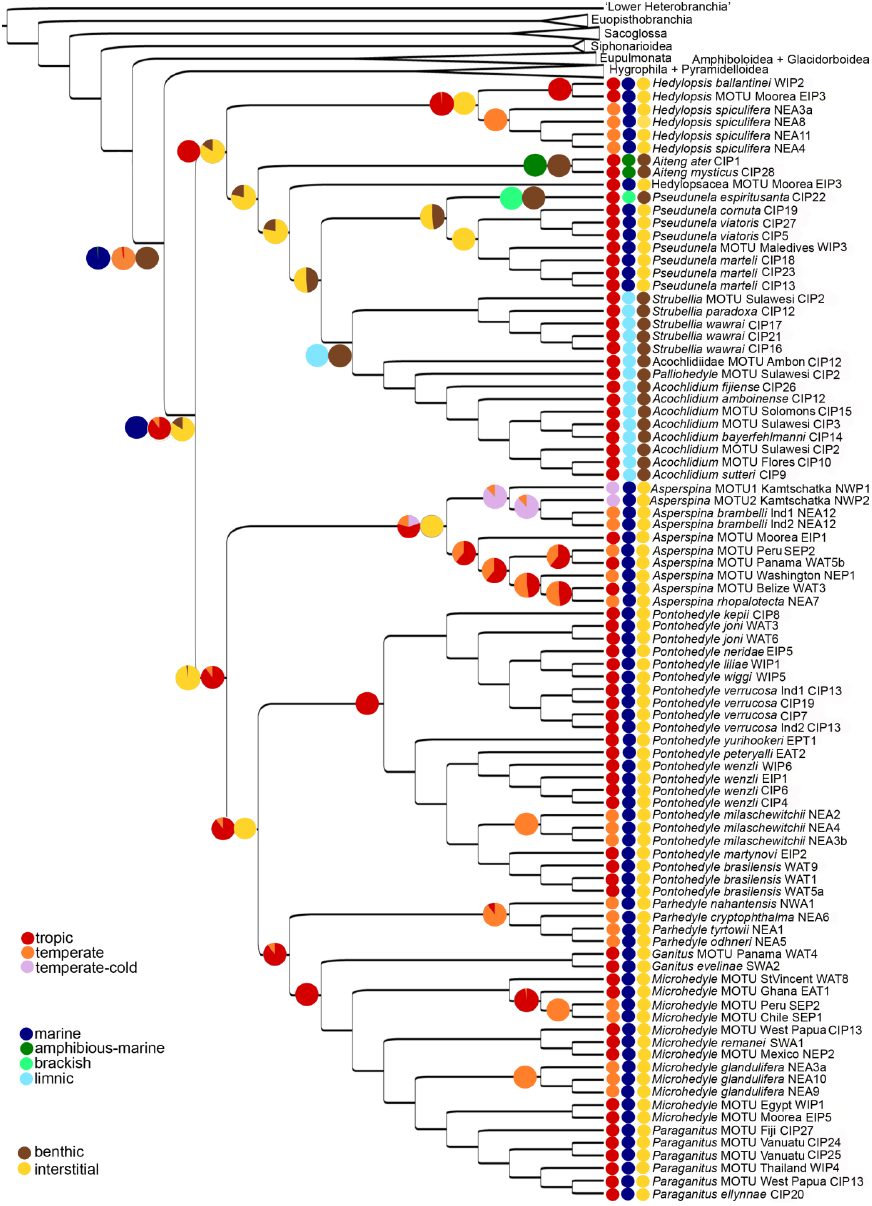
Ancestral state reconstruction of ecological traits (climate, habitat and life style) in Acochlidia, retrieved in Mesquite on the maximum-likelihood phylogeny shown in Fig. 3, outgroups collapsed.

The origin of the hedylopsacean radiation clearly occurred in tropical waters (lh 0.99), with *Hedylopsis spiculifera* currently being the only representative found in temperate waters. The radiation of the hedylopsacean clade shows a remarkable flexibility in habitat choice: the hedylopsacean ancestor likely inhabited the marine (lh0.99) mesopsammon (lh 0.84, most parsimoniously benthic or interstitial) and shifts to a benthic life style occurred in parallel three times: 1) along the stemline of Aitengidae with a shift to amphibious-marine habitat, 2) within Pseudunelidae with a shift to brackish habitat, and 3) at the base of Acochlidiidae with the invasion of the limnic system. Alternatively (but with weaker statistical support lh 0.21), the ancestor of Aitengidae+(Hedylopsacea sp.+(Pseudunelidae+Acochlidiidae)) resumed a benthic lifestyle with two independent invasions back into the mesopsammon in Hedylopsacea sp. and Pseudunelidae. Within Pseudunelidae+Achochlidiidae hypotheses on the ancestral lifestyle are unclear: either the common ancestor was 1) benthic (lh 0.48) and within Pseudunelidae, the slugs are secondarily mesopsammic or 2) mesopsammic (lh 0.52) and the transition to a benthic lifestyle occurred twice independently (within Acochlidiidae and for *Pseudunela espiritusanta*). Testing the alternative topology suggested by Bayesian inference (see Fig. 4) with Aitengidae forming a basal offshoot to the remaining Hedylopsacean clades only slightly weakens the likelihood for an interstitial hedylopsacean ancestor (lh 0.78), but does not affect the results on ancestral character states described above and shown in Fig. 6. The entire diversification of Microhedylacea occurred in the marine mesopsammon, shifts are restricted to climatic zones, originating in tropical waters and inhabiting temperate zones at least five times independently (Fig. 6). *Asperspina* is the only acochlidian clade with members occurring in temperate-cold waters, but there is only a low probability that the asperspinid ancestor already inhabited temperate (lh 0.22) or temperate-cold waters (lh 0.20). Adaptation to cold waters probably (lh 0.88) occurred along the (*A. brambelli*+*Asperspina* MOTU Kamtschatka) stemline and between temperate and tropic waters another three times independently (Fig. 3). Additionally, the radiation of the genus *Parhedyle* occurred in temperate waters (lh 0.89) and it is currently the only genus with no representatives known from tropical waters. The ancestor of *Pontohedyle* inhabited tropical waters. *Pontohedyle milaschewitchii* is the only lineage in the tropical *Pontohedyle* clade, which secondarily adapted to temperate waters. Within paraphyletic *Microhedyle* an adaptation to temperate waters occurred twice independently in the Eastern Pacific and the Northern Atlantic.

### 4 Discussion

#### 4.1 Acochlidian origin and phylogeny

The exact position of Acochlidia within panpulmonate euthyneurans remains unresolved in this study. Schrödl and Neusser (2010) demonstrated that parsimony analyses of morpho-anatomical characters easily lead to a grouping of Acochlidia with other minute, mesopsammic taxa as an artifact of multiple convergent adaptations. This explained previous phylogenetic hypotheses that are incompatible with recent molecular results, such as the idea of a common origin with Rhodopemorpha (Salvini-Plawen and Steiner, 1996) or equally minute Runcinacea and Cephalaspidea (Wägele and Klussmann-Kolb, 2005). Molecular analyses have considerably rearranged the classical view on euthyneuran phylogeny, commonly rejecting monophyly of ‘Opisthobranchia’ and ‘Pulmonata’ and placing the traditional opisthobranch order Acochlidia in (pan)pulmonate relationships (Jörger et al., 2010b; Klussmann-Kolb et al., 2008). ‘Opisthobranchia’ are also in conflict with morphological analyses (Dayrat and Tillier, 2002; Haszprunar, 1985; Wägele et al., 2014; Wägele et al., 2008). Revisiting morphological characters of Euthyneura (*sensu* Jörger et al. (2010b)) in the light of the new phylogenetic hypothesis, the presence of rhinophores innervated by N3 (*nervus rhinophoralis*) might be an apomorphy for Euthyneura (Brenzinger et al., 2013; Wägele et al., 2014) and a gizzard (i.e. muscular oesophagial crop lined with cuticula) apomorphic for Euopisthobranchia (Jörger et al., 2010b). Currently, the double rooted rhinophoral nerve or the double rooted procerebrum presents the only putative morphological apomorphy for the highly diverse panpulmonate clade including Acochlidia (Brenzinger et al., 2013). Molecular studies on euthyneuran phylogeny provided high bootstrap support for most higher panpulmonate taxa (Dayrat et al., 2011; Dinapoli and Klussmann-Kolb, 2010; Jörger et al., 2010b; Klussmann-Kolb et al., 2008), but fail to recover decent support values on the relationships among major clades within Panpulmonata.

Acochlidia are suggested to be sister to (Pyramidelloidea+Amphiboloidea)+Eupulmonata (Klussmann-Kolb et al., 2008) or Eupulmonata (Jörger et al., 2010b). In comparison to these previous studies the outgroup sampling herein was optimized and targeted to include representatives of all major lineages among the putative panpulmonate relatives. Despite this denser taxon sampling we again fail to reconstruct sister group relationships of Acochlidia reliably, i.e. with significant support. In the present analyses Acochlidia are sister to a clade comprised of Hygrophila+(Pyramidelloidea+(Amphiboloidea+Glacidorboidea)), all together sister to Eupulmonata (see Fig. 2). Our analyses indicate an explosive radiation of panpulmonate diversity (expressed by very short branches at the base of the higher taxa, see Figs. 3, 4). The chosen ‘standard marker’ set (i.e. partial mitochondrial COI and 16S rRNA and nuclear 18S rRNA and 28S rRNA), offers the broadest taxon sampling currently available for any molecular markers in Mollusca (Stöger et al., 2013), but unfortunately at the present stage seems to be incapable of solving the deep panpulmonate relationships reliably. This is probably handicapped by the old Mesozoic origin of Panpulmonata in combination with a very rapid diversification into the higher taxa in the Mesozoic Jurassic (see Fig. 4 and Additional material 3). The application of new molecular markers with potential phylogenetic signal for deep euthyneuran splits, such as phylogenomic datasets (e.g., Kocot et al., 2013), but at the same time allowing for a similarly broad and representative taxon sampling as gathered using the ‘standard markers’, are overdue.

In contrast to ambiguous sister group relationships, the monophyly of Acochlidia is currently undisputed based on morphology (Schrödl and Neusser, 2010) and molecular markers (present study; Jörger et al., 2010b). Even though the two major acochlidian clades – Hedylopsacea and Microhedylacea – differ remarkably in their evolutionary history (see discussion below), they share a unique combination of characters (e.g. characteristic separation into head-foot complex and visceral hump, presence of calcareous spicules and a pre-pharyngeal nerve ring with separated cerebral and pleural ganglia and a rhinophoral nerve innervating the rhinophore) (Schrödl and Neusser, 2010). Many of these features can be interpreted as either plesiomorphic in the new heterobranch tree or apomorphic through progenetic reductions, i.e. result of the ‘meiofaunal syndrome’ (Brenzinger et al., 2013). The recently discovered Aitengidae (Swennen and Buatip, 2009), which were placed into hedylopsacean Acochlidia based on microanatomy and molecular data, conflict with the classical acochlidian bauplan by lacking the division into fully retractable head-foot complex and visceral hump (Neusser et al., 2011a). Our present phylogenetic hypothesis implies that Aitengidae have lost this most striking acochlidian apomorphy secondarily, as several of the features have been reduced secondarily in other genera as well (e.g. rhinophores in microhedylacean *Pontohedyle* and *Ganitus*). The monophyly of Acochlidia was again recovered in our study independent of the type of analyses (maximum-likelihood see Fig. 3 or Bayesian interference see Fig. 4). Even though the monophyly was not well supported – probably suffering from the same effects as discussed above for Panpulmonata in general – there is currently no doubt that Acochlidia are monophyletic.

The first detailed classification of Acochlidia conducted by Rankin (1979), which led to the establishment of 19 genera in 13 families and four suborders for only 25 nominal species, was heavily criticized by all subsequent authors (Arnaud et al., 1986; Schrödl and Neusser, 2010; Wawra, 1987). The alternative classification erected by Wawra (1987) rendered the system of Rankin (1979) obsolete and was largely confirmed by first cladistic analyses of Acochlidia relying on over a hundred biological and morphological characters (Schrödl and Neusser, 2010). Our molecular phylogenetic hypothesis (see Fig. 3) is largely compatible with the one based on morphological markers (Schrödl and Neusser, 2010, see fig.3). Therefore, most of the potential synapomorphies hypothesized for higher acochlidian clades and major evolutionary scenarios are confirmed, such as the progressive elaboration of copulatory organs within hedylopsaceans (Neusser et al., 2009a). Also the apomorphic, successive reduction of reproductive organs among microhedylaceans is supported, with loss of copulatory organs in the common ancestor of Asperspinidae plus Microhedylidae (s.l.), and evolution of secondary gonochorism in the latter (e.g., Schrödl and Neusser, 2010; Sommerfeldt and Schrödl, 2005). Unfortunately, our analyses lack monotypic Tantulidae. Thorough 3D-microanatomy from the type material showed mixed and partially unique characters states for *Tantulum* (e.g. complex, unarmed anterior copulatory complex), which justifies the erection of a monotypic family by Rankin (1979), based on its unique ecology and morphological features (Neusser and Schrödl, 2007). Cladistic analyses (Schrödl and Neusser, 2010) based on morphological and biological characters resolved *Tantulum* as basal offshoot of Hedylopsacea, i.e. sister to (Hedylopsidae+(Pseudunelidae+Acochlidiidae)). Due to additional inclusion of Aitengidae and the undescribed Hedylopsacea sp. into Hedylopsacea, *Tantulum* cannot be plotted unambiguously onto our molecular tree any longer. Tantulidae retains several supposedly plesiomorphic features (e.g. an unarmed copulatory organ) and likely diverges quite early among Hedylopsacea; it probably originates in the late Mesozoic (i.e. Cretaceous) (see basal Hedylopsacean diversification in Fig. 4).

All of the included acochlidian families recognized by Schrödl and Neusser (2010) were monophyletic, largely with high bootstrap supports (see Fig. 3). The only disagreement to morphological approaches is paraphyletic Ganitidae (*Ganitus*+*Paraganitus*). Ganitidae which receives high support in morphological analyses, is nested here within paraphyletic *Microhedyle,* and was thus included in a broader definition of Microhedylidae *s.l.* (Schrödl and Neusser, 2010). Within Microhedylacea, only few distinguishing characters are present, leaving most of the clade unresolved in morphological analyses (see Schrödl and Neusser, 2010). The monophyly of Ganitidae is supported mainly by the common characteristics of the dagger-shaped radula and the related pharyngeal characteristics (i.e. paired cuticular mandibles) (Schrödl and Neusser, 2010). The paired cuticular mandibles are discussed to serve as functional ‘odontophore’ and counterparts for the muscles moving the radula (Challis, 1968; Marcus, 1953). Gosliner (1994) originally suspected a relationship of Ganitidae with Sacoglossa, which possess a strikingly similar dagger-shaped radula, used for piercing algae. All currently available morphological and molecular data on Ganitidae clearly reject a sacoglossan relationship of this family and places it rather in a derived microhedylacean relationship (present study; Jörger et al., 2010b; Neusser et al., 2011a; Schrödl and Neusser, 2010). Thus, radula similarities seem to be based on functional convergence rather than common ancestry, supported by considerable differences in pharyngeal morphology (i.e. presence of mandibles, longitudinal pharynx musculature and absence of an ascus in Ganitidae) (B. Eder, pers. comm.; Challis, 1968; Marcus, 1953). Based on our molecular data, the dagger-shaped radulae and remarkably similar pharyngeal structures (B. Eder, pers. comm.) of *Ganitus* and *Paraganitus* also refer to convergent modifications potentially based on a shared feeding strategy in the mesopsammon.

In contrast to morphological analyses, our study successfully resolved the Microhedylidae s.l. confirming monophyletic *Pontohedyle*, *Parhedyle*, *Ganitus* and *Paraganitus*, but rendering *Microhedyle* paraphyletic. The paraphyly of *Microhedyle* is concordant with morphological analyses, but due to the high degree of homoplasy and few morphological apomorphies supporting the genera, we are currently unable to diagnose the different clades within *Microhedyle* based on morphological characters and thus leave ‘*Microhedyle*’ for a future taxonomic revision of Acochlidia. As discussed previously, *Microhedyle nahantensis* and *M. odhneri* need to be transferred to *Parhedyle* based on molecular data and the presence of a special type of asymmetric radulae (formula 1.1.2, with inner right lateral tooth smaller than outer one) (Eder et al., 2011, own unpublished data), diagnostic for the genus *Parhedyle* (Wawra, 1987).

Additionally, our study detected 30 additional acochlidian lineages, which could not be assigned directly to valid species, and were determined as molecular operational taxonomic units (MOTUs) herein. All MOTUs were independent lineages in phylogenetic analyses, but only some were supported by unique morphological characters recognizable in the field and via light microscopy (e.g. Hedylopsacea sp. or Acochlidiidae sp., own unpublished data). Recent studies have shown that acochlidian diversity can best be tackled by using an integrative taxonomic approach making use of all available character sets, e.g. microanatomy, DNA sequence data and biogeography (Jörger et al., 2012; Neusser et al., 2011b). The presented MOTUs, thus, still require comparative microanatomical approaches with their sister groups, a molecular species delineation approach capable of dealing with low numbers of specimens, as described by Jörger et al. (2012), and molecular diagnoses (see Jörger and Schrödl, 2013) in addition to morphological ones, if any exist. This is beyond the scope of the present paper.

#### 4.2 Timeframe and biogeography of acochlidian evolution

##### The Mesozoic origin

Previous molecular clock analyses on euthyneuran gastropods indicated a rapid radiation into the modern higher taxa in the early to mid Mesozoic (Jörger et al., 2010b; Klussmann-Kolb et al., 2008; Stöger et al., 2013). Our present molecular clock estimates – relying on the same set of genetic markers – supported these previous diversification estimates. The variation in fossil calibration points and the stability throughout our sensitivity analyses herein (see Additional material 2) shows that the estimations do not rely on a single calibration point but correspond well to a broad set of euthyneuran fossils. Earlier terrestrial gastropods fossils from the Paleozoic interpreted as pulmonates (Solem, 1985; Solem and Yochelson, 1979) are contradicted by our time trees (Additional material 3). Pulmonate affinities of such fossils also would imply long gaps in the fossil record. Our results thus support earlier criticism (Dayrat et al., 2011; Mordan and Wade, 2008) that suggests the few detectable characters of such potential Palaeozoic pulmonate shells might as well refer to prosobranch lineages. Considerably earlier molecular clock estimates dating the radiation of euthyneurans to the Cambrian and the appearance of (pan)pulmonate lineages to Ordovician and Silurian (Medina et al., 2011) based on mitogenomic data was likely driven by erroneous coding procedures and biased sampling (Schrödl et al., 2011b), or that mitogenomic datasets available to date are simply not suitable to resolve basal euthyneuran topologies correctly (Stöger and Schrödl, 2013).

The rapid mid Mesozoic radiation of euthyneuran and especially panpulmonate taxa (and thus the origin of Acochlidia) might be an outcome of the breakup of the supercontinent Pangaea and Panthalassas Ocean (approx. 180 mya) and resultant isolation of marine biotas, which in general boosted marine diversity (Cox and Moore, 2010; Lomolino et al., 2010). Based on this hypothesis (which relies on molecular clock data), the origination of the major panpulmonate clades including Acochlidia might be characterized as part of the successful recovery fauna of the Triassic/Jurassic turnover.

For gastropod taxa with a Mesozoic origin and poor fossil record, precise areas of origin are difficult to determine and ancestral area reconstructions based on molecular phylogenies could often only support assumptions of origins in the Tethyan realm, without further specification (Frey and Vermeij, 2008; Malaquias and Reid, 2009; Ozawa et al., 2009; Reid et al., 2010). As shown in previous studies on Mesozoic radiations for other taxa, Acochlidia originated when shelf areas of oceans varied dramatically from the current profiles and it is thus difficult to assign recent distribution ranges to historic areas. Based on current knowledge, Acochlidia are limited to rather shallow shelf areas; we thus aimed to allocate recent continental margins to the historic topologies at the ages of the reconstructed nodes and discuss the biogeography accordingly. While there is strong support that the ancestral acochlid inhabited the marine mesopsammon in tropical waters (see Fig. 6), none of our ancestral area reconstructions (DEC-model and S-DIVA) provided reliable support values for the ancestral area of Acochlidia (see Fig. 4). The most likely ancestral areas for Acochlidia based on the DEC model is the North-Eastern part of the Tethys Ocean and the Western margin of the Panthalassic/Pacific Ocean, but only with marginal higher relative probabilities than other regions of the Tethys or the forming Atlantic Ocean. Although our data clearly indicated a circum-tropical distribution of Acochlidia throughout the Tethys, Pacific and Atlantic Ocean (see Discussion below), the precise region of origin could not be solved. The evolution of Microhedylacea is characterized by numerous dispersal, vicariance and local extinction events (see Fig. 4). The indicated local extinction events might either truly relate to extinction events in the corresponding areas or refer to still unsampled microhedylacean lineages in the different areas of the world’s oceans. The complex picture on microhedylacean biogeography with sister clades in non-connected areas of distribution, and somewhat long internal branches, demonstrates the high degree of missing diversity with the current ancestral chronogram only able to present current knowledge. In the present study we included almost 85 % of the described acochlidian species-level diversity and added approximately another 50 % by putatively new species (left as undetermined MOTUs for future research). Nevertheless, our sampling map (see Fig. 1) shows ‘white spots’ of large biogeographic areas, which are still relatively unexplored for meiofaunal molluscs. In particular, the Western Indo-Pacific (former Western Tethys Region) which was revealed as a major area of transition and dispersal for many microhedylacean lineages (see Fig. 4) and discussed as center of origin and diversification in the Oligocene and early Miocene for marine molluscan biodiversity (Harzhauser et al., 2007) is virtually unsampled, as is the vast majority of the Eastern Atlantic. Based on their patchy occurrence and with large parts of tropical sands still unsampled, we are probably only scraping at the surface of the expected recent acochlidian diversity, which still remains to be discovered. Evidently, this might have major influence on biogeographical hypotheses, but our robust phylogenetic hypothesis – concordant between previous morphological (Schrödl and Neusser, 2010) and molecular analyses (Jörger et al., 2010b) – nevertheless, allows for the reconstruction of ancestral area chronograms and the reflection how historic geographic events triggered acochlidian diversification:

##### Distributing circumtropical and beyond

A warm-tropical circum-equatorial seaway (Tethyan Seaway) formed during the early Cretaceous (Cox and Moore, 2010; Lomolino et al., 2010), and throughout the Cretaceous and Paleogene it served as gateway for marine biotas between the tropical world’s oceans (Frey and Vermeij, 2008). This connectivity is well reflected in our dataset on Acochlidia by numerous dispersal events across the Tethys Ocean, the forming Atlantic and East Pacific Ocean (see Fig. 4). The Tethyan Seaway is described as a strong westward directed current, providing better dispersal conditions westward than eastward (Cox and Moore, 2010). This is not supported, however, by our data, which shows frequent changes in the direction of dispersal events even within a clade (see e.g. *Pontohedyle* and *Asperspina* in Fig. 4). The current abated once the Tethyan Seaway narrowed by the convergence of Africa with Eurasia in Oligocene and Miocene, enabling an eastward dispersal from the East Pacific to the Western Atlantic (Lomolino et al., 2010). The independent dispersal events in Asperspinidae eastwards via the tropical East Pacific into the Caribbean Sea predate the narrowing and closure of the Tethyan Seaway, estimated to the early Paleogene, but the dispersal into the Atlantic might indeed have occurred after the weakening of the circum-tropical current. Mesopsammic Acochlidia are considered as poor dispersers due to the lack of a planktonic larval stage (veliger larvae remain in the interstices of sand grains (Swedmark, 1968)). Thus, currents at local scales, the presence of continuous coastlines and the uplift of ocean floors creating shallow shelf seas or terranes, providing stepping stones for dispersal probably had larger influence on their historic biogeography than major oceanic current systems in the past.

In the Mesozoic, terranes in the central Pacific are thought to have served as stepping stones which allowed dispersal into the Eastern Indo-Pacific (Grigg and Hey, 1992). However, our data presented five independent dispersal events in Hedylopsacea and Microhedylacea from the CIP to EIP (the island of Moorea) not limited to the Mesozoic, but instead ranging between Upper Cretaceous and Late Miocene (88–7 mya). If we are not severely underestimating acochlidian dispersal abilities, this indicates that remote Eastern Indo-West Pacific islands were continuously accessible either via island stepping stones or seamounts.

The East Pacific Barrier (EPB, approx. 5000km of deep water separating the Indo-West Pacific fauna from the tropical East Pacific fauna) is today generally considered as one of the most effective barriers to the dispersal of marine shallow-water fauna and data from fossil coral suggests that it was largely in place throughout the Cenozoic (Grigg and Hey, 1992). Based on our global phylogeny the ancestors of *Pontohedyle* (Microhedylidae *s. l.*) crossed the EPB once in an eastward dispersal event (approx. 82 mya) and once westward (approx. 54 mya). The ancestor of *Asperspina* MOTU Moorea also dispersed in westward direction from the Northeast Pacific (approx. 63 mya) (see Fig. 4). Reports on gene exchange or closely related species spanning the EPB (e.g., in gastropod species with long larval stages (Reid et al., 2010), echinoderms (Lessios et al., 1998) and tropical fish (Lessios and Robertson, 2006)) indicate that this potent barrier is not ‘impassable’. The span of the EPB by a presumably poor disperser like *Pontohedyle* or *Asperspina*, however, is still puzzling. All acochlidian dispersal events dated to Upper Cretaceous till late Paleocene/early Eocene, and fossil data on coral suggest that the EPB was less effective during the Cretaceous (Grigg and Hey, 1992). The long stemlines of the discussed species, however, might also indicate that the evolutionary history of these clades is not well covered by our dataset and might lack e.g. Western Atlantic sister clades which could reverse the dispersal picture.

Asperspinidae were reconstructed with an ancestral range spanning the North Pacific and presented the only clade with a putative trans-arctic dispersal in the late Paleogene into NEA, which is in concordance with the usual dispersal direction of most other Mollusca in the Trans-Artic Interchange between the two northern Oceans (Vermeij, 1991). Asperspinidae also show the highest flexibility to ocean temperatures and include the only currently known polar acochlid *Asperspina murmanica* (see Kudinskaya and Minichev, 1978; Neusser et al., 2009b), which unfortunately could not be included in the present analyses. The tropical origin of Asperspinidae received high statistical support in our analyses and the cold water adaptation likely evolved at the base of the NWP/NEA clade in the late Eocene (see Figs. 3, 4). This correlates to a shift of the warm-temperate zones southward in the circumpolar region following a series of temperature declines after the mid Eocene and the establishment of a cold-temperate regime (Briggs, 2003).

##### Major vicariance events shaping the evolutionary history of Acochlidia

Our molecular clock estimates, the pan-Tethyan, Central American and Pacific distribution of acochlidian taxa and the supposed poor dispersal abilities of Acochlidia, all suggest that the biogeography of the clade is shaped by major vicariance events during the Mesozoic and Cenozoic: the closure of the Tethyan Seaway (Tethyan Terminal Event - TTE) in the Miocene (approx. 18 mya), collision of Australia and New Guinea with Eustralasia forming the modern Wallacea province (25 mya) and the Pliocene uplift of the Isthmus of Panama (3.4 resp. 2.1 mya) (Cox and Moore, 2010; Lomolino et al., 2010).

The Terminal Tethyan Event in the Miocene (TTE, i.e. the vicariance event preventing exchange between the Atlantic and Indo-Pacific regions) (Cox and Moore, 2010; Lomolino et al., 2010) is reflected in our data in the split between Mediterranean and North East Atlantic *Microhedyle glandulifera* and its sister clade - the (central) Indo-West Pacific radiation of *Paraganitus*. The split was estimated slightly prior to the TTE by our molecular clock analyses (approximately 40–18 mya, see Fig. 4 and Additional material 3). Data from other gastropod species pairs, which presumably originated by Tethyan vicariance, however, also predate the final closure of the seaway in the Miocene (e.g., Malaquias and Reid, 2009; Reid et al., 2010), probably related to the proceeding isolation of the Tethyan realm prior to the final closure. Assuming a Tethyan divergence via vicariance between *M. glandulifera* and *Paraganitus*, one would expect an eastward dispersal pattern in the historic biogeography of *Paraganitus*, which rather presents a mixed picture with westward tendencies (see Fig. 4).

Nowadays, the central Indo-West Pacific forms a hotspot in marine biodiversity, but based on fossil records of Mollusca the faunal diversity was rather poor until the late Paleogene (Frey and Vermeij, 2008). During Oligocene times the Western Tethys region potentially served as center of origin and diversification of molluscan taxa, which shifted southeast after the TTE (Harzhauser et al., 2007). Our data showed that 1) hedylopsacean Aitengidae, Pseudunelidae and Acochlidiidae evolved *in situ* in the central Indo-Pacific Ocean and 2) no *in situ* splits within the central Indo-West Pacific occurred prior to the mid Paleogene. The estimated timeframe of the origin of the families predates, however, the closure of Tethyan Seaway but no data on relict species in other parts of the world suggest an origin elsewhere. The late Oligocene and Miocene is known for being a time in which the diversity of marine shallow-water fauna increased, likely influenced by the availability of new shallow-water habitats formed by the collision of Australia and New Guinea with the southeastern edge of Eurasia (Williams and Duda, 2008). Williams and Duda (2008) showed increased rate of cladogenesis in different unrelated gastropod genera in the Indo-West Pacific in the beginning of late Oligocene/early Miocene. This is in concert with the increased diversification especially in hedylopsacean Indo-West Pacific clades starting in the late Oligocene, which likely holds major responsibility for the detected overall increase in diversification rates of Acochlidia in the Oligocene by the birth-death-shift model (see Fig. 5B).

Moreover, our data showed two recent specification events by vicariance in the Indo-West Pacific: in hedylopsacean *Pseudunela marteli* (CIP) and *Pseudunela* MOTU Maledives (WIP), and microhedylacean *Paraganitus* from CIP and *Paraganitus* MOTU Thailand (WIP). Strong currents in recent central Indo-West Pacific waters should facilitate the dispersal of larvae and be responsible for the invisibility of the Wallace’s Line (sharp transition between terrestrial faunas of eastern and western Indonesia) in marine taxa (Barber et al., 2000). But Barber et al. (2000) showed that Pleistocene ocean basins are still reflected in the genetic structure of shrimp populations, which might resemble relicts of Indian and Pacific populations separated by the emergence of the Sunda and Sahul continental shelves, suggesting the presence of a marine Wallace’s line perpendicular to the terrestrial Wallace’s line. Our data by far predates the Pleistocene Indonesian continental shelf formations, but might represent relicts reflecting the better connectivity between Indian and Pacific populations prior to tectonic events in the Wallacea region in the early Neogene. An integrative species delineation approach, however, still needs to confirm the species status of the identified MOTUs; a putative conspecificity between the different populations would imply genetic exchange between the recent CIP and WIP populations, thus contradicting assumptions on a marine Wallace’s line, while a subdivision of MOTUs would further support it.

Following the TTE the North Eastern Atlantic, viz. Mediterranean suffered from an impoverishment in faunal diversity, due to the Miocene cooling, Messinian salinity crisis and glacial events in late Pliocene and Pleistocene (Harzhauser et al., 2007). Although our data recovered NEA frequently as ancestral area of microhedylacean clades throughout the Mesozoic and Paleogene, only comparably few lineages persisted or still radiated in NEA after the TTE. The only entirely North Atlantic clade *Parhedyle* originated in the NEA prior to the closure if the Tethyan Seaway and dispersed westward into the NWA, following the typical unidirectional expansion of molluscan species across (or around) the Atlantic (Vermeij, 2005). Our molecular clock estimate on the radiation of NEA *Parhedyle* (approx. mya) slightly predated the Messinian salinity crisis, but in absence of records of these *Parhedyle* species outside the Mediterranean, it is likely that the radiation occurred in the Mediterranean during reinvasion after the crisis.

Based on our data, Western Atlantic species of Acochlidia presented relictual Tethys origins, inhabiting the area prior to the TTE (in *Pontohedyle*) or showed origins in the New World (WAT+EPT) prior to the closure of the Isthmus of Panama (in *Asperspina*) (see Fig. 4). *Ganitus* and western Atlantic *Microhedyle* radiated in the Western Atlantic. Two ‘soft barriers’ for dispersal in marine taxa in the Atlantic (Briggs and Bowen, 2013) are reflected in our data, the vicariance event splitting eastern and western Atlantic *Microhedyle* and *Asperspina* species (open water expanse of the mid-Atlantic) and the vicariance event splitting the two *Ganitus* MOTUs, which according to our time tree slightly predated the freshwater discharge of the Amazon river (11 mya, see Briggs and Bowen, 2013). In a previous study on microhedylid *Pontohedyle* population this soft barrier between Caribbean and Brazilian provinces, however, still allows for gene exchange (Jörger et al., 2012). Vicariance events potentially related to the formation of the Isthmus of Panama (approx. 3 mya, see Cox and Moore, 2010) are reflected twice within our dataset: *Asperspina* MOTU Peru (SEP) and *Asperspina* MOTU Panama (WAT) and *Microhedyle remanei* (SWA, but also reported from WAT) and *Microhedyle* MOTU Mexico (NEP), even though this implies a series of still unsampled (or extinct) intermediate populations.

##### Evolutionary hotspots

To summarize the discussed biogeography with regard to evolutionary hotspots of Acochlidia, our reconstructions of ancestral ecological traits like climate (Fig. 6) clearly suggest that the tropical regions provided the species pools of Acochlidia, from which temperate regions were populated several times independently. In concert with data on other gastropod taxa (see e.g., Williams, 2007; Williams and Duda, 2008), Acochlidia show a latitudinal gradient in taxonomic diversity and diversification concentrated in the tropics (see Figs. 1, 6). Generalizing, the Indo-West Pacific – especially its central region – is remarkably diverse in comparison to marine geographic regions of the Eastern Pacific or Atlantic, with the Western Atlantic showing an intermediate diversity of marine taxa and the Eastern Atlantic and Pacific harboring the lowest diversity (Briggs, 2007). The vast majority of worldwide mesopsammic fauna is still unexplored, but our data already offers some indication on the centers of acochlidian evolution. The tropical Western Atlantic is equally well-sampled as the central Indo-Pacific (concerning number of stations), however, only one enigmatic hedylopsacean species (*Tantulum elegans*) has been reported, while the central Indo-West Pacific harbors the vast majority of hedylopsacean lineages and acts as cradle for the recent Cenozoic diversification. Hypothetically, including *Tantulum* into our ancestral area chronogram, late Mesozoic Hedylopsacea probably had a circum-tethyal distribution. One can speculate that *Tantulum* presents a relict in an unstable geological area, which was majorly affected by the mass extinction events like the Cretaceous-Paleogene (approx. 66 mya) or Eocene-Oligocene event (approx. 33 mya). Fossil data suggests that up to 90 % of the molluscan fauna was eliminated in Gulf of Mexico during each event (Hansen et al., 2004); similar ranges might be expected for the Caribbean Sea, but none of the events can be traced on our diversification rate estimates (see Fig. 5). Since no marine hedylopsacean has yet been discovered in the Western Atlantic, no rough dating or speculations on the evolutionary background of the shift from marine to freshwater in Tantulidae can be done. *Tantulum* remains truly enigmatic as the only minute, interstitial or mud-dwelling slug in freshwater (Neusser and Schrödl, 2007; Rankin, 1979). While the sluggish bauplan in freshwater is unique to Acochlidia in general, all other known acochlidian limnic slugs reestablished a benthic lifestyle.

#### 4.3 Habitat shifts

##### Into the mesopsammon

The interstitial or mesopsammic fauna is one of the most diverse on Earth and comprises minute representatives from most major lineages of Metazoa, including several exclusively meiofaunal clades (Rundell and Leander, 2010; Worsaae et al., 2012). The physical parameters of the interstitial habitat of marine sands are challenging for the inhabitants, but likely persisted throughout the history of Eukaryota, and a meiofaunal lifestyle might have evolved even prior to the Cambrian with numerous independent colonization events since then (Rundell and Leander, 2010). A previous study based on molecular analyses discussed at least five independent interstitial invasions within heterobranch slugs and dated colonization events range between the Mesozoic and Cenozoic (Jörger et al., 2010b). Based on our data herein the invasion of the mesopsammon already occurred along the stemline of Acochlidia and could thus be estimated to Lower to Middle Jurassic (see Fig. 4, Additional material 3).

The restricted space in the interstitial habitat requires great morphological and biological adaptations for its minutely-sized inhabitants. Even though the exact origin of Acochlidia within panpulmonates remain unresolved, no evidence exists at present that any of the sister groups are derived from meiofaunal ancestors, thus it is likely that the miniaturization of Acochlidia occurred along with the transition into the mesopsammon in the acochlidian ancestor. Many meiofaunal taxa show paedomorphic traits (i.e. morphological characters present in juveniles or larvae of closely related species) discussed as result of progenesis (accelerated sexual maturation in relation to somatic development) or neoteny (retardation of somatic development in relation to sexual maturity) (Brenzinger et al., 2013; Rundell and Leander, 2010; Westheide, 1987; Worsaae et al., 2008; Worsaae et al., 2012). Little is still known of the influence of progenesis in the evolution of meiofaunal slugs, mainly due to a lack of ontogenetic data and uncertain phylogenetic affinities. Within the basal heterobranch Rhodopemorpha, strong evidence for a progenetic origin is reported, based on paedomorphic traits in adults including features like the pentaganglionate stage of the visceral loop and a protonephridial organization of the kidney (Brenzinger et al., 2013). Jörger et al. (2010b) suggests a progenetic origin of the acochlidian ancestor by relating the typical acochlidian morphology (shell-less yet free visceral hump) to the phenotype of an abnormally developed larva reported in the nudibranch *Aeolidiella alderi* and some stylommatophoran Eupulmonata (Tardy, 1970), leading to a very similar external morphology.

Life in the interstitial spaces of sand grains evidently constrains morphology and leads to a series of convergent adaptations in the different taxa (e.g., vermiform body shape), termed the ‘meiofaunal syndrome’ (Brenzinger et al., 2013). Our study confirms assumptions based on morphological data that the ancestral acochlid had already invaded the interstitial habitat, which was related to major morphological adaptations (e.g. minute, worm-shaped body, loss of shell, detorsion resulting in symmetric body condition) (Schrödl and Neusser, 2010). The ancestral acochlid probably possessed a simple sac-like kidney that is typical for marine Euthyneura (Neusser et al., 2011b), and had a hermaphroditic, phallic, monaulic reproductive system, transferring sperm via copulation (Schrödl and Neusser, 2010). Miniaturized body plans often combine simplified organ systems with morphological novelties and inventions (Hanken and Wake, 1993; Westheide, 1987). The microhedylacean clade lacks novel morphological inventions entirely, and rather present a line of ‘regressive evolution’ (i.e. as simplification and reduction of organ systems (Swedmark, 1964, 1968)). All microhedylacean Acochlidia studied in sufficient detail (Eder et al., 2011; Jörger et al., 2008; Jörger et al., 2007; Neusser et al., 2006; Neusser et al., 2009b) have a simple sac-like kidney, as described as ancestral for Acochlidia (Neusser et al., 2011b). So far no habitat transitions are known for Microhedylacea. Most members of the clade are limited to truly marine and subtidal sands, but minor tolerance to changes in salinity concentration seem to be present in e.g. in the microhedylacean species reported from the Black Sea *P. milaschewitchii* and *Parhedyle tyrtowii* (Kowalevsky, 1901). Few microhedylacean species (e.g. *Asperspina murmanica, A. riseri and Pontohedyle verrucosa*) occur in the intertidal, which is characterized by temporary changes in salinity either due to rain or desiccation. No specific adaptations of the excretory or circulatory system, however, are reported in these slugs (Challis, 1970; Kowalevsky, 1901; Morse, 1976; Neusser et al., 2009b) and thus do not seem to be mandatory to deal with the temporary osmotic stress in the intertidal.

In regards to the reproductive system, some hedylopsacean Acochlidia are protandric hermaphrodites and the ontogenetic loss of male genital organs was interpreted as the precursor for the evolution of secondary gonochorism at the base of Microhedylidae s.l. (Schrödl and Neusser, 2010), which might be related to a second progenetic event in acochlidian evolution. Microhedylacea are characterized by the loss of copulatory organs and the usage of spermatophores for sperm transfer (Jörger et al., 2009; Schrödl and Neusser, 2010). The male genital opening is usually located dextrolaterally or shifted anteriorly and spermatophores are randomly attached to the mates followed by dermal insemination (Jörger et al., 2009; Morse, 1994; Swedmark, 1968). Spermatophores are a common development across different meiofaunal taxa and considered as adaptation to the spatially restricted and unstable interstitial habitat, favoring imprecise but fast sperm transfer (Jörger et al., 2009). What serves as advantageous mode of reproduction in the mesopsammic world might present an evolutionary disadvantage, however, in transition to freshwater or semi-terrestrial habitats where directly injected sperm via stylets or regular copulation is more common. From the origin of Microhedylidae s.l in the mid-Jurassic (approx. 166 mya), this family remained in almost complete morphological stasis: minor changes occurred only in form, shape and number of spicules, the secondary reduction of the rhinophores (twice independently in *Pontohedyle* and *Ganitus*) and modifications of the radula (i.e. small second right lateral tooth in *Parhedyle* and twice independently the evolution of a dagger-shaped radula with corresponding modifications of the pharynx in *Ganitus* and *Paraganitus*). Other than that, due to the reduced stage of all organ systems, even advanced microanatomy failed to recover distinguishing features even at the genus level (for details on the anatomy see e.g. Eder et al., 2011; Jörger et al., 2010a; Jörger et al., 2008; Jörger et al., 2007; Neusser et al., 2006). Moreover, intraspecific or even intraindividual (right and left side) variation, especially within the nervous system, often exceeds interspecific morphological variation in these highly cryptic lineages. This high intraspecific variability is discussed as consequence from miniaturization and paedomorphosis, frequently involving late forming structures, which are then individually truncated in development (Hanken and Wake, 1993). The most striking example of morphological stasis in Microhedylacea is a recently discovered world-wide radiation in *Pontohedyle* slugs which present clearly independently evolving genetic lineages but are entirely cryptic based on traditional taxonomic characters (i.e. external morphology, spicules and radula features). Even microanatomy does not reveal reliable diagnostic characters (Jörger et al., 2012).

In direct contrast to the morphological reduction and simplification in Microhedylacea, the evolution of Hedylopsacea is driven by novel inventions and increasing organ complexity. The development of a complex kidney at the base of Hedylopsacea probably plays a key role for the ecological diversification of the clade, serving as precursor to habitat shifts which enables to handle osmotic stress (Brenzinger et al., 2011b; Neusser et al., 2009a; Neusser et al., 2011b; Neusser and Schrödl, 2009).

##### Out of the mesopsammon

The evolutionary pathway away from the meiofaunal lifestyle is even less studied and understood than the way in. Meiofaunal animals may have played a major role in the earliest diversification of bilaterians (Rundell and Leander, 2010; Worsaae et al., 2012), but few reports document the reversion of miniaturized forms re-establishing a benthic life-style and evolving a ‘secondary gigantism’ in body size (see e.g., Westheide, 1982; Worsaae and Kristensen, 2005). In an example known for annelids, morphological features which were reduced as previous adaptation to the interstitial life (e.g. setae) were not reestablished with increasing body size (Westheide, 1982). Our ancestral state reconstruction of Acochlidia supports a mesopsammic hedylopsacean ancestor and three independent ways out of the interstitial habitat and to larger body sizes: in Aitengidae, *Pseudunela espiritusanta* and Acochlidiidae (see Fig. 6). Alternatively, but less supported in the analyses, the benthic lifestyle could have been reestablished only once (or twice) in an ancestral lineage within Hedylopsacea, then again Pseudunelidae (and Hedylopsacea sp.) would have secondarily recolonized the interstitial habitat. Currently, all transitions out of the mesopsammon are always into habitats characterized by non-marine salinities (brackish, limnic, amphibious) (see Fig. 6). In absence of any macrofaunal marine Hedylopsacea, the ‘secondary gigantism’ seems a consequence of the habitat transition into brackish water, freshwater and terrestrial systems rather than the necessary precursor of the habitat switch e.g., to be able to cope with osmoregulatory requirements. The hypothesis, that habitat shifts primarily depend on the ability of efficient osmoregulation independent of the overall body size is further supported by the presence of the limnic, yet minute and interstitial *Tantulum elegans*. Some recently discovered deep-sea benthic slugs, however, show putative hedylopsacean relationships (TPN unpublished data) and might shed new light on hedylopsacean habitat shifts out of the mesopsammon in the future.

##### Shifting from marine to (semi-)terrestrial and limnic habitats

Evolutionary transitions between aquatic and terrestrial habitats as well as between marine and freshwater systems are comparatively rare in the animal kingdom (Vermeij and Dudley, 2000). Therefore considerable barriers probably exist among the habitats related to osmoregulation, desiccation and novel predator-prey interactions (Vermeij and Dudley, 2000; Vermeij and Wesselingh, 2002). The key to the diversification of particularly panpulmonate Euthyneura was the invasion of freshwater and terrestrial habitats, and the drivers of those habitat transitions are of major interest (Barker, 2001; Klussmann-Kolb et al., 2008). Earlier assumptions that pulmonate taxa secondarily invaded the marine habitat via terrestrial pathways (Solem, 1985) are nowadays replaced by a consensus on the marine origin of (Pan)Pulmonata (Dayrat et al., 2011; Klussmann-Kolb et al., 2008). This is confirmed by our ancestral state reconstruction, in which most deep panpulmonate nodes were highly supported as marine (lh 0.96 -1.0, reconstruction not shown). Truly terrestrial panpulmonates are found within Stylommatophora, Systellommatophora and Ellobioidea and several independent evolutionary pathways led to the life on land in pulmonate slugs and snails (Barker, 2001; Dayrat et al., 2011), probably via marine marginal zones and amphibious-marine transition stages (Klussmann-Kolb et al., 2008). Klussmann-Kolb et al. (2008) suggested that the colonization of freshwater in Euthyneura was a unique evolutionary event (in Hygrophila) directly from marine habitat via an aquatic pathway. Our ancestral area reconstruction confirmed the proposed aquatic pathway into freshwater in Hygrophila and also for Glacidorboidea, ‘amphibious-limnic’ lineages in Ellobioidea and limnic Acochlidia. This also indicates that conclusions on a unique transition event into freshwater, however, were premature and do not reflect the highly complex picture of habitat transitions in Panpulmonata.

Because of incomplete taxon sampling, conclusions from ancestral area reconstructions across these phylogenies have to be carefully evaluated. Although it is ideal to all major lineages of each outgroup clade, ideally with a basal rather than derived internal position, the selected taxa may not necessarily reflect the ecological variability and basal state of habitat within their clade, e.g., the non-inclusion of the putatively basal marine *Williamia* within Siphonarioidea resulted in the amphibious semi-terrestrial ancestral state for the diversification of the clade (results not shown). Due to the uncertainty of panpulmonate sister group relationships and the incomplete ecological representation of our panpulmonate outgroups (e.g., Siphonarioidea, Hygrophila and Ellobioidea) we refrain from reporting ancestral characters states across Panpulmonata in detail, including dating ancestral areas for the involved transitions. Nevertheless, our present and previous molecular clock analyses (Jörger et al., 2010b; Klussmann-Kolb et al., 2008) indicate that some of the major habitat transitions (e.g., land invasion by the stylommatophoran ancestor and shift to freshwater by the ancestor of Hygrophila) date back to the Mesozoic. In contrast, our study on Acochlidia revealed comparably recent habitat transitions within Hedylopsacea: According to our analyses, freshwater was invaded only once in the ancestor of Acochlidiidae (see Fig. 6). Another, independent transition to freshwater occurred in the Western Atlantic Tantulidae (Schrödl and Neusser, 2010), a taxon unfortunately unavailable for molecular approaches (see Discussion above). The semi-terrestrial habitat was invaded once in Aitengidae and permanently brackish water once at the base of Pseudunelidae by *Pseudunela espiritusanta* (see Fig. 6). All habitat shifts occurred in the central Indo-West Pacific. The habitat shifts to limnic and brackish water were dated herein to the late Paleogene. Unfortunately, the stemline of Aitengidae spans nearly 130 my hindering assumptions on the timeframe in which the transition to a (semi-) terrestrial lifestyle occurred, but it is likely that they fall in the same timeframe in which the forming Wallacea region offered ideal conditions for shifts in habitat. During the Paleogene the Central Indo-West Pacific region was under major geological reformation and lots of new shallow shelf areas and islands appeared (Cox and Moore, 2010; Lomolino et al., 2010). This 1) provided comparatively pristine habitats with low levels of competition and predation for newcomers, which is discussed as beneficial for habitat shifts (Vermeij and Dudley, 2000) and 2) boosted marine diversity in general, thus raising the competition and predation pressure in the old habitat. Based on our phylogenetic hypotheses the (semi-)terrestrial habitat in Aitengidae was invaded by a marine ancestor and can be considered part of the marginal zone between marine and terrestrial habitat. With regard to the colonization of limnic habitats however, paticular hypotheses of brackish habitats serving as stepping stones into freshwater ones are not supported by our phylogeny and the reconstruction of ancestral character states, including those on temporarily brackish habitats (e.g., intertidal mesopsammic *Pseudunela cornuta* influenced by rain) and permanently brackish waters (e.g., *P. espiritusanta*) (Neusser et al., 2009a; Neusser and Schrödl, 2009). But more intermediate taxa inhabiting zones with decreasing salinity are necessary to reconstruct the evolutionary scenario of the invasion of limnic systems in Acochlidia.

#### 4.4 Morphological and behavioral adaptations

Habitat shifts require adaptation to the new physical environment; to deal with osmotic stress, risk of desiccation and adapt life strategies concerning reproduction, predator avoidance and available food sources (Mordan and Wade, 2008). In the evolution of Hedylopsacea, the complexity of the excretory and circulatory systems increased in taxa that conquered new habitats, compared with their fully marine, mesopsammic sister groups. Next to ‘secondary gigantism’, which improves the volume/surface ratio for osmoregulation, the excretory system of limnic and brackish Acochlidiidae and Pseudunelidae potentially increases effectiveness by evolving a long, looped nephroduct (which is short in fully marine sister species of Pseudunelidae) (Neusser et al., 2009a; Neusser et al., 2011b; Neusser and Schrödl, 2009). ‘Enhanced’ nephroducts are known for limnic panpulmonates (Smith and Stanisic, 1998), where it is long and looping as in some hedylopsacean Acochlidia (e.g. Hygrophila, *Acroloxus* and *Ancylus* (Delhaye and Bouillon, 1972)) or bladder-like (*Glacidorbis* (Ponder, 1986)). The nephroduct is also elaborate in terrestrial Systell- and Stylommatophora, but not modified in the remaining coastal or amphibious panpulmonate taxa (e.g. Siphonariidae (Delhaye and Bouillon, 1972) or sacoglossan *Gascoignella* (Kohnert et al., 2013)). Additionally, specialized heart cells in limnic *Strubellia*, which probably enhance circulation, likely present another novel adaptive feature to life in freshwater (Brenzinger et al., 2011b). The dorsal vessel system (modified part of the kidney) in Aitengidae, was proposed as an adaptation to a semi-terrestrial lifestyle, potentially used for oxygen supply in a gill-less animal (Neusser et al., 2011a).

Analogous to the increasing complexity in excretory and circulatory systems, several ‘novel’ or reinvented reproductive features also evolved in Hedylopsacea. The Acochlidiidae developed a progressively large and complex copulatory apparatus, with additional glands and cuticular injection systems, culminating in the ‘giant’ phallus equipped with several rows of cuticular spines termed ‘rapto-penis’ (Schrödl and Neusser, 2010). In addition, sperm storage organs such as a seminal receptacle and a bursa copulatrix (gametolytic gland) are absent in basal acochlidians, but reappear in derived hedylopsaceans (Schrödl and Neusser, 2010). On one hand, this increasing diversity and complexity of male and female reproductive organs, paired with an obviously traumatic type of mating (Lange et al., 2013), suggests an arms race between individuals and perhaps even sexes in these hermaphrodites. Large reproductive organs are coupled with secondarily large body sizes and a benthic lifestyle. On the other hand, this re-establishment of reproductive features present in euthyneuran outgroups but putatively reduced in basal mesopsammic acochlidian lineages, shows that the ontogenetic source of these structures might persist and can be reactivated in larger individuals again. This implies, however, that the function of theses developmental pathways is maintained by other selective constrains, otherwise a reactivation after such long timescales is considered impossible (Marshall et al., 1994).

The mode of reproduction is further modified in limnic Acochliididae, which produce a large number of eggs in contrast to the low reproductive output in the marine sister taxa. Benthic, limnic acochlidiids are likely amphidromic, i.e. have a marine planktonic larval stage and recolonize freshwater as juveniles (Brenzinger et al., 2011b). The freshwater fauna of tropical oceanic islands in the central Indo-West Pacific is dominated by amphidromous species (Crandall et al., 2010). This life strategy evolved several times independently among invertebrates, likely because it facilitates (re-)invasion of island habitats and unstable stream environments (Crandall et al., 2010; McDowall, 2007). This long-distance marine planktonic dispersive stage promotes a population structure similar to those in marine species with planktonic larvae, and is characterized by little genetic structure among the populations of different island archipelagos (Crandall et al., 2010; Kano and Kase, 2004). The dispersal abilities of the unique ‘adhesive larvae’ reported for Acochlidiidae are unknown and it remains speculative whether other animals are used as dispersal vectors or whether the glue is used on the substrate to avoid being swamped to the open sea (Brenzinger et al., 2011b). In the future population genetic analyses might give insights about the connectivity of populations on different islands and thus allow conclusions on the fate of the larvae after being washed into the sea. It is likely that these changes in habitat and mode of reproduction result in entirely different population structures, with rather well-connected populations and widespread species in limnic taxa versus a high degree of endemism and small distributions in marine, mesopsammic Acochlidia.

The pathway into the mesopsammon is thought to be advantageous in avoiding predator pressure (Palmer, 1988). In general, slugs should be more vulnerable to predation than snails, as the gastropod shell serves as protection against predators. However, sea slug lineages which lack a protective shell have independently evolved a series of defensive mechanisms such a cleptocnides, acid glands or secondary metabolites (for summary see Wägele and Klussmann-Kolb, 2005). In Acochlidia, no special defensive features are known apart from their unique ability to retract their head-foot complex into their visceral hump (Schrödl and Neusser, 2010), showing similar behavior to snails withdrawing into their shell. The acochlidian visceral hump can be equipped with a dense arrangement of calcareous spicules in Asperspinidae and Hedylopsidae forming a secondary ‘spicule shell’ (Schrödl and Neusser, 2010; Sommerfeldt and Schrödl, 2005). The ability to retract the anterior is present in all mesopsammic Acochlidia, and may be an effective defense against (small-sized) meiofaunal predators by increasing the body diameter to avoid being swallowed whole, and additionally protects essential body parts against bites. Secondarily benthic Acochlidia can only slightly or partially retract, which may result from the changes in overall morphology (e.g., the flattened leaf-like visceral sac in *Acochlidium* and *Palliohedyle*). But probably the strategy is not particularly efficient against large-sized benthic predators such as crabs or fish and might have lost its evolutionary significance in benthic environments. Interestingly, no marine benthic acochlidians are described at present, thus acochlidian evolution lacks evidence for the most direct habitat shift (from the marine mesopsammon back to a benthic marine lifestyle, see discussion above). The transitions to a benthic lifestyle in Acochlidia occur where habitats present comparatively low predator pressure, such as freshwater systems, or by dwelling in brackish water like *Pseudunela espiritusanta* (Neusser and Schrödl, 2009), and in the latter, is supported by behavioral predator avoidance such as hiding beneath stones during daytime. The semi-terrestrial Aitengidae is an exception, and lives in the marine intertidal habitat which is very stressful due to highly fluctuating temperatures and salinities, the risk of desiccation and the presence of many predators such as crabs, other arthorpods or sea birds. Behavioral stress avoidance, like hiding into the damp crevices of intertidal rocks during daytime in *Aiteng mysticus* (Neusser et al., 2011a), might have been the key to successfully colonize this habitat. Whether or not Aitengidae possess additional defensive features, for example via chemical substances, still needs to be explored in future research.

Given the late Paleogene timeframe and the ancestral area (Wallacea) for the major habitat shifts in Acochlidia, the driving forces for the transitions still remain unclear. Food sources of Acochlidia are largely unknown, but it is likely that they are highly specialized feeders, e.g., preying on the eggs of co-occurring species. It can be speculated that the availability of new food sources with less competition might play a role, e.g. *Aiteng ater*, which specializes on feeding on insect pupae (Swennen and Buatip, 2009). Moreover, potential co-evolution between limnic Acochlidia like *Strubellia* and the neritid gastropods whose eggs they feed on (Brenzinger et al., 2011b), should be investigated in future research combining gut content analyses, molecular clock analyses and the habitat shifts in both groups.

## Conclusions

The Acochlidia provide an astonishing example of two major evolutionary clades differing enormously in habitat transition flexibility. The hedylopsacean evolution presents a mosaic of habitat transitions between aquatic and (semi-) terrestrial habitats shifting from the mesopsammon to epibenthic lifestyle and vice versa, which corresponds to a series of novel morphological developments and increasing complexity e.g., in excretory and reproductive features within some hedylopsacean clades. Consequently, Hedylopsacea comprise high morphological plasticity and ecological diversity with their major diversity hotspot in the central Indo-West Pacific, and morphological divergent lineages are still expected in further research. Conversely, their sister clade Microhedylacea remained in almost entire morphological and ecological stasis since the late Mesozoic. Miniaturization, organ simplifications and specialization led to a highly successful clade, which is worldwide distributed with several transitions to temperate and even temperate-cold waters, occurs in locally high species densities and has apparently successfully survived or recolonized areas after major extinction events. But this evolutionary success of Microhedylacea by taking the ‘regressive’ and specialized pathway into the mesopsammon apparently also forms a dead-end road concerning morphological or ecological diversification. The currently known Microhedylacean diversity is largely cryptic, which is also to be expected for their still undiscovered lineages. Adding about 30 new MOTUs, the present study again confirms the existence of hidden marine diversity, and highlights the more general need for integrative species delimitation and the potential for molecular description of cryptic species.

The comparatively small and well-studied clade of Acochlidia demonstrates the high degree of habitat flexibility in panpulmonates, evoking potential complexity in the evolutionary history of other less-known panpulmonate clades. The present study shows how habitat transitions can be placed in space, time and biological context, once a robust, integrative species-level phylogeny is established, and despite limitations such as still undiscovered diversity. Even though panpulmonate relationships cannot be satisfactorily resolved at present, converging molecular clock data from sensitivity analyses with different fossil calibration points already indicate that different shifts in habitat in Panpulmonata occurred in different Mesozoic and Cenozoic timeframes and therefore various geological and ecological backgrounds. The ancestral marine habitat of the basal panpulmonate Acochlidia and other deep panpulmonate nodes is the originator of the panpulmonate evolution in the mid Mesozoic, but habitat shifts need to be addressed individually across each clade.

## Acknowledgements

We want to thank our collaborators for sharing their material on Acochlidia with us, without their contributions the present study would be worse of several important lineages. The following people have contributed material and/ or helped to arrange sampling permits: Fontje Kaligis and Gustav Mamangkey (for material from Indonesia), Yuri Hooker (for support in Peru), Peter Ryall (for support in Ghana), Greg Rouse (for material from Moorea), the organizing team of the World Congress for Malacology 2010 (for sampling permits in Thailand), the Red Sea Environmental Center (for support in collecting material in Egypt), Tanya Korshunova (for material from Russia), Kevin Kocot (for material from the US West Coast), the Dumbarton Agricultural Station (for permits in St. Vincent), and Jon Norenburg, Katrine Worsaae, Rick Hochberg and other participants of the Encyclopedia of Life Meiofauna Workshop (for sorting material in the Caribbean). A special thanks goes to Yasunori Kano for material from Palau and valuable discussions acochlidian evolution and habitat shifts. The SANTO 2006 Expedition to Vanuatu was organized by Museum national d’Histoire naturelle, Paris, Pro-Natura International (PNI), and Institut de Recherche pour le Développement (IRD). It operated under a permit granted to Philippe Bouchet (MNHN) by the Environment Unit of the Government of Vanuatu. The Marine Biodiversity part of the expedition, apart from the Census of Marine Life’s CReefs program, was specifically funded by grants from the Total Foundation and the Sloan Foundation. Tanja Stadler is thanked for her patient support in TreePar analyses and Prashant Sharma and Richard Ree for their help with Lagrange. Warm thanks go to Katrine Worsaae for inspiring discussions on the evolution of meiofauna. This study was supported by a PhD scholarship of the Volkswagen Foundation to KJ and the DFG grant SCHR667/4 to MS. AVM received funding by the Russian Foundation for Basic Research (grant # 13-04-01641a).

## Additional material

### Additional material 1

Sampling localities of Acochlidia included in the present study (recollecting attempts at the same position are marked with a and b). Localities referring to type localities of valid acochlidian species are marked with *.

Collectors: AA – Andreas Altenöder, PB – Pat Boaden, LD – Ludwig Demharter, AD – Angela Dinapoli, BE – Barbara Eder, GH – Gerhard Haszprunar, MH – Martin Heß, KJ – Katharina Jörger, YK – Yasunori Kano, KK – Kevin Kocot, AM – Alexander Martynov, RM – Roland Meyer, TN – Timea Neusser, GR – Greg Rouse, JS – Julia Sigwart, MS – Michael Schrödl, ES – Enrico Schwabe, NW – Nerida Wilson

### Additional material 2

Summary of the different molecular clock analyses performed in this study (and compared to Jörger et al. 2010) and the resulting estimated node ages for the major acochlidian taxa.

### Additional material 3

Complete chronogram of Acochlidia, panpulmonate outgroups shown. Divergence times obtained from BEAST v1.6.1 under a relaxed uncorrelated clock model, node ages indicated at nodes, bars represent 95 % highest posterior densities (only presented for nodes with a PP > 0.5).

## References

Altschul, S.F., Gish, W., Miller, W., Myers, E.W., Lipman, D.J., 1990. Basic local alignment search tool. J Mol Biol 215, 403–410.

Arnaud, P.M., Poizat, C., Salvini-Plawen, L.v., 1986. Marine-interstitial Gastropoda (including one freshwater interstitial species). In: Botosaneanu, L. (Ed.), Stygofauna Mundi. Brill/Backhuys, Leiden, pp. 153–161.

Bandel, K., 1994. Triassic Euthyneura (Gastropoda) from St. Cassian Formation (Italian Alps) with a discussion on the evolution of the Heterostropha. Freib Forschh 2, 79–100.

Barber, P.H., Palumbi, S.R., Erdmann, M.V., Moosa, M.K., 2000. Biogeography - A marine Wallace’s line? Nature 406, 692–693.

Barker, G.M., 2001. Gastropods on land: Phylogeny, diversity and adaptive morphology. In: Barker, G.M. (Ed.), The biology of terrestrial molluscs. CAB International, Oxon, New York, pp. 1–146.

Bernt, M., Bleidorn, C., Braband, A., Dambach, J., Donath, A., Fritzsch, G., Golombek, A., Hadrys, H., Juhling, F., Meusemann, K., Middendorf, M., Misof, B., Perseke, M., Podsiadlowski, L., von Reumont, B., Schierwater, B., Schlegel, M., Schrödl, M., Simon, S., Stadler, P.F., Stöger, I., Struck, T.H., 2013. A comprehensive analysis of bilaterian mitochondrial genomes and phylogeny. Mol Phylogenet Evol 69, 352–364.

Brenzinger, B., Haszprunar, G., Schrödl, M., 2013. At the limits of a successful body plan – 3D microanatomy, histology and evolution of *Helminthope* (Mollusca: Heterobranchia: Rhodopemorpha), the most worm-like gastropod. Front Zool 10, 37.

Brenzinger, B., Neusser, T.P., Glaubrecht, M., Haszprunar, G., Schrödl, M., 2011a. Redescription and three-dimensional reconstruction of the limnic acochlidian gastropod *Strubellia paradoxa* (Strubell, 1892) (Gastropoda: Euthyneura) from Ambon, Indonesia. J Nat Hist 45, 183–209.

Brenzinger, B., Neusser, T.P., Jörger, K.M., Schrödl, M., 2011b. Integrating 3D microanatomy and molecules: Natural history of the Pacific freshwater slug *Strubellia* Odhner, 1937 (Heterobranchia, Acochlidia), with description of a new species. J Molluscan Stud 77, 351–374.

Briggs, J.C., 2003. Marine centres of origin as evolutionary engines. J Biogeogr 30, 1–18.

Briggs, J.C., 2007. Marine longitudinal biodiversity: causes and conservation. Divers Distrib 13, 544–555.

Briggs, J.C., Bowen, B.W., 2013. Marine shelf habitat: biogeography and evolution. J Biogeogr 40, 1023-1035.

Bücking, G., 1933. *Hedyle amboinensis* (Strubell). Zool Jb Syst 64 549–582.

Challis, D.A., 1968. A new genus and species of the order Acochlidiacea (Mollusca: Opisthobranchia) from Melanesia. Trans R Soc NZ Zool 10, 191–197.

Challis, D.A., 1970. *Hedylopsis cornuta* and *Microhedyle verrucosa*, two new Acochlidiacea (Mollusca: Opisthobranchia) from the Solomon Islands Protectorate. Trans R Soc NZ 12, 29–40.

Cox, C.B., Moore, P.D., 2010. The engines of the planet I: Plate tectonics. In: Cox, C.B., Moore, P.D. (Eds.), Biogeography. An ecological and evolutionary approach. John Wiley and Sons Inc., pp. 153–173.

Crandall, E.D., Taffel, J.R., Barber, P.H., 2010. High gene flow due to pelagic larval dispersal among South Pacific archipelagos in two amphidromous gastropods (Neritomorpha: Neritidae). Heredity 104, 563–572.

Dayrat, B., Conrad, M., Balayan, S., White, T.R., Albrecht, C., Golding, R., Gomes, S.R., Harasewych, M.G., de Frias Martins, A.M., 2011. Phylogenetic relationships and evolution of pulmonate gastropods (Mollusca): New insights from increased taxon sampling. Mol Phylogenet Evol 59, 425–437.

Dayrat, B., Tillier, S., 2002. Evolutionary relationships of euthyneuran gastropods (Mollusca): a cladistic re-evaluation of morphological characters. Zool J Linn Soc 135, 403–470.

Delhaye, W., Bouillon, J., 1972. L’évolution et l’adaptation de l’organe excréteur chez les Mollusques gastéropodes pulmonés. I. Introduction générale et histophysiologie comparée du rein chez les Basommatophores. Bull Biol France Belg 106, 46–77.

Dinapoli, A., Klussmann-Kolb, A., 2010. The long way to diversity - Phylogeny and evolution of the Heterobranchia (Mollusca: Gastropoda). Mol Phylogenet Evol 55, 60–76.

Dinapoli, A., Zinssmeister, C., Klussmann-Kolb, A., 2011. New insights into the phylogeny of the Pyramidellidae (Gastropoda). J Molluscan Stud 77, 1–7.

Drummond, A., Ashton, B., Buxton, S., Cheung, M., Cooper, A., Heled, J., Kearse, M., Moir, R., Stones-Havas, S., Strurrock, S., Thierer, T., Wilson, A., 2010. Geneious v5.4. www.geneious.com.

Drummond, A.J., Rambaut, A., 2007. BEAST: Bayesian evolutionary analysis by sampling trees. BMC Evol Biol 7.

Eder, B., Schrödl, M., Jörger, K.M., 2011. Systematics and redescription of the european meiofaunal slug *Microhedyle glandulifera* (Kowalevsky, 1901) (Heterobranchia: Acochlidia): evidence from molecules and morphology. J Molluscan Stud 77, 388–400.

Frey, M.A., Vermeij, G.J., 2008. Molecular phylogenies and historical biogeography of a circumtropical group of gastropods (Genus: *Nerita*): Implications for regional diversity patterns in the marine tropics. Mol Phylogenet Evol 48, 1067-1086.

Gosliner, T.M., 1994. Gastropoda: Opisthobranchia. In: Harrison, F.W., Kohn, A.J. (Eds.), Microscopic anatomy of invertebrates, Mollusca I. Wiley-Liss, Ltd., pp. 253–355.

Grande, C., Templado, J., Cervera, J.L., Zardoya, R., 2004. Molecular phylogeny of the Euthyneura (Mollusca: Gastropoda). Mol Biol Evol 21, 303–313.

Grigg, R.W., Hey, R., 1992. Paleoceanography of the tropical Eastern Pacific Ocean. Science 255, 172–178.

Hall, T.A., 1999. BioEdit: a user-friendly biological sequence alignment editor and analysis program for Windows 95/98/NT. Nucleic Acids Symp Ser 41, 95–98.

Hanken, J., Wake, D.B., 1993. Miniturization of body size - organismal and evolutionary significance. Annu Rev Ecol Syst 24, 501–519.

Hansen, T.A., Kelley, P.H., Haasl, D.M., 2004. Paleoecological patterns in molluscan extinctions and recoveries: comparison of the Cretaceous-Paleogene and Eocene-Oligocene extinctions in North America. Palaeogeogr Palaeoclimatol Palaeoecol 214, 233–242.

Harrison, A.D., Rankin, J.J., 1976. Hydrobiological studies of Easter Lesser Antillean Islands I. St. Vincent: Freshwater fauna - its distribution, tropical river zonation and biogeography. Arch Hydrobiol 50, 275–311.

Harzhauser, M., Kroh, A., Mandic, O., Piller, W.E., Gohlich, U., Reuter, M., Berning, B., 2007. Biogeographic responses to geodynamics: A key study all around the Oligo-Miocene Tethyan Seaway. Zool Anz 246, 241–256.

Haszprunar, G., 1985. The Heterobranchia - a new concept of the phylogeny of the higher Gastropoda. Z Zool Syst Evolutionsforsch 23, 15–37.

Haynes, A., Kenchington, W., 1991. *Acochlidium fijiensis* sp. nov. (Gastropoda: Opisthobranchia: Acochlidiacea) from Fiji. Veliger 34, 166–171.

Jörger, K.M., Heß, M., Neusser, T.P., Schrödl, M., 2009. Sex in the beach: spermatophores, dermal insemination and 3D sperm ultrastructure of the aphallic mesopsammic *Pontohedyle milaschewitchii* (Acochlidia, Opisthobranchia, Gastropoda). Mar Biol 156, 1159–1170.

Jörger, K.M., Kristof, A., Klussmann-Kolb, A., Schrödl, M., 2010a. Redescription of the meiofaunal gastropod *Parhedyle cryptophthalma*, with focus on nervous system and sensory organs. Spixiana 33, 161–170.

Jörger, K.M., Neusser, T.P., Brenzinger, B., Schrödl, M., 2014. Exploring the diversity of mesopsammic gastropods: How to collect, identify, and delimitate small and elusive sea slugs? Am Malacol Bull 32, 290–307.

Jörger, K.M., Neusser, T.P., Haszprunar, G., Schrödl, M., 2008. Undersized and underestimated: 3D-visualization of the Mediterranean interstitial acochlidian gastropod *Pontohedyle milaschewitchii* (Kowalevsky, 1901). Org Divers Evol 8, 194–214.

Jörger, K.M., Neusser, T.P., Schrödl, M., 2007. Re-description of a female *Pontohedyle brasilensis* (Rankin, 1979), a junior synonym of the Mediterranean *P. milaschewitchii* (Kowalevsky, 1901) (Acochlidia, Gastropoda). Bonn Zool Beitr 55, 283–290.

Jörger, K.M., Norenburg, J.L., Wilson, N.G., Schrödl, M., 2012. Barcoding against a paradox? Combined molecular species delineations reveal multiple cryptic lineages in elusive meiofaunal sea slugs. BMC Evol Biol 12, 245.

Jörger, K.M., Schrödl, M., 2013. How to describe a cryptic species? Practical challenges of molecular taxonomy. Front Zool 10, 59.

Jörger, K.M., Stöger, I., Kano, Y., Fukuda, H., Knebelsberger, T., Schrödl, M., 2010b. On the origin of Acochlidia and other enigmatic euthyneuran gastropods, with implications for the systematics of Heterobranchia. BMC Evol Biol 10, 323.

Kano, Y., Kase, T., 2004. Genetic exchange between anchialine cave populations by means of larval dispersal: the case of a new gastropod species *Neritilia cavernicola*. Zool Scr 33, 423–437.

Katoh, K., Kuma, K., Toh, H., Miyata, T., 2005. MAFFT version 5: improvement in accuracy of multiple sequence alignment. Nucleic Acids Res 33, 511–518.

Klussmann-Kolb, A., Dinapoli, A., Kuhn, K., Streit, B., Albrecht, C., 2008. From sea to land and beyond-new insights into the evolution of euthyneuran Gastropoda (Mollusca). BMC Evol Biol 8, 57.

Kocot, K.M., Halanych, K.M., Krug, P.J., 2013. Phylogenomics supports Panpulmonata: Opisthobranch paraphyly and key evolutionary steps in a major radiation of gastropod molluscs. Mol Phylogenet Evol.

Kohnert, P., Brenzinger, B., Jensen, K., Schrödl, M., 2013. 3D-microanatomy of the semiterrestrial slug *Gascoignella aprica* Jensen, 1985—a basal plakobranchacean sacoglossan (Gastropoda, Panpulmonata). Org Divers Evol 13, 583–603.

Kohnert, P., Neusser, T.P., Jörger, K.M., Schrödl, M., 2011. Time for sex change! 3D-reconstruction of the copulatory system of the ‘aphallic’ *Hedylopsis ballantinei* (Gastropoda, Acochlidia). Thalassas 27, 113–119.

Kowalevsky, A., 1901. Les Hédylidés, étude anatomique. Mem Acad Imp Sci St-Petersbourg 12, 1–32.

Kudinskaya, E.V., Minichev, Y.S., 1978. Psammological essays. I. The organization and systematic position of the mollusc *Hedylopsis murmanica* n. sp. (Opisthobranchia, Acochlidiida). Trudy Petergofskogo Biologicheskogo Instituta Leningradskogo Gosudarstvennogo Universiteta 26, 69–86.

Lange, R., Reinhardt, K., Michiels, N.K., Anthes, N., 2013. Functions, diversity, and evolution of traumatic mating. Biol Rev (Camb) 88, 585–601.

Lessios, H.A., Kessing, B.D., Robertson, D.R., 1998. Massive gene flow across the world’s most potent marine biogeographic barrier. Proceedings of the Royal Society B: Biological Sciences 265, 583–588.

Lessios, H.A., Robertson, D.R., 2006. Crossing the impassable: genetic connections in 20 reef fishes across the eastern Pacific barrier. Proceedings of the Royal Society B: Biological Sciences 273, 2201–2208.

Lomolino, M.V., Riddle, B.R., Whittaker, R.J., Brown, J.H., 2010. Biogeography. Sinauer Associates, Sunderland, MA, USA.

Maddison, W.P., Maddison, D.R., 2011. Mesquite: a modular system for evolutionary analysis. Version 2.75. In: http://mesquiteproject.org (Ed.).

Malaquias, M.A.E., Reid, D.G., 2009. Tethyan vicariance, relictualism and speciation: evidence from a global molecular phylogeny of the opisthobranch genus *Bulla*. J Biogeogr 36, 1760–1777.

Marcus, E., 1953. Three Brazilian sand-Opisthobranchia. Bol Fac Filos Ci Letr Univ Sao Paulo, Zool 164, 165–203.

Marshall, C.R., Raff, E.C., Raff, R.A., 1994. Dollo’s law and the death and resurrection of genes. Proceedings of the National Academy of Sciences 91, 12283-12287.

McDowall, R.M., 2007. On amphidromy, a distinct form of diadromy in aquatic organisms. Fish Fish 8, 1–13.

Medina, M., Lal, S., Vallès, Y., Takaoka, T.L., Dayrat, B.A., Boore, J.L., Gosliner, T., 2011. Crawling through time: Transition of snails to slugs dating back to the Paleozoic, based on mitochondrial phylogenomics. Marine Genomics 4, 51–59.

Meier, R., Shiyang, K., Vaidya, G., Ng, P.K.L., 2006. DNA barcoding and taxonomy in diptera: A tale of high intraspecific variability and low identification success. Syst Biol 55, 715–728.

Misof, B., Misof, K., 2009. A monte carlo approach successfully identifies randomness in multiple sequence alignments: A more objective means of data exclusion. Syst Biol 58, 21–34.

Mordan, P.B., Wade, C.M., 2008. Heterobranchia II: The Pulmonata. In: Ponder, W.F., Lindberg, D.R. (Eds.), Phylogeny and evolution of the Mollusca. University California Press, Berkeley Los Angeles London, pp. 409–426.

Morse, M.P., 1976. *Hedylopsis riseri* sp. n., a new interstitial mollusc from the New England Coast (Opisthobranchia, Acochlidiacea). Zool Scr 5, 221–229.

Morse, M.P., 1994. Current knowledge of reproductive biology in two taxa of interstitial molluscs (class Gastropoda: order Acochlidiacea and class Aplacophora: order Neomeniomorpha). In: Wilson, W.H., Stricker, S.A., Shinn, G.L. (Eds.), Reproduction and development of marine invertebrates. John Hopkins University Press, pp. 195–205.

Neusser, T.P., Fukuda, H., Jörger, K.M., Kano, Y., Schrödl, M., 2011a. Sacoglossa or Acochlidia? 3D reconstruction, molecular phylogeny and evolution of Aitengidae (Gastropoda: Heterobranchia). J Molluscan Stud 77, 332–350.

Neusser, T.P., Heß, M., Haszprunar, G., Schrödl, M., 2006. Computer-based three-dimensional reconstruction of the anatomy of *Microhedyle remanei* (Marcus, 1953), an interstitial acochlidian gastropod from Bermuda. J Morphol 267, 231–247.

Neusser, T.P., Heß, M., Schrödl, M., 2009a. Tiny but complex - interactive 3D visualization of the interstitial acochlidian gastropod *Pseudunela cornuta* (Challis, 1970). Front Zool 6, 20.

Neusser, T.P., Jörger, K.M., Schrödl, M., 2011b. Cryptic species in tropic sands - Interactive 3D anatomy, molecular phylogeny and evolution of meiofaunal Pseudunelidae (Gastropoda, Acochlidia). PLoS ONE 6, e23313.

Neusser, T.P., Martynov, A.V., Schrödl, M., 2009b. Heartless and primitive? 3D reconstruction of the polar acochlidian gastropod *Asperspina murmanica*. Acta Zool (Stock) 90, 228–245.

Neusser, T.P., Schrödl, M., 2007. *Tantulum elegans* reloaded: a computer-based 3D-visualization of the anatomy of a Caribbean freshwater acochlidian gastropod. Invertebr Biol 126, 18–39.

Neusser, T.P., Schrödl, M., 2009. Between Vanuatu tides: 3D anatomical reconstruction of a new brackish water acochlidian gastropod from Espiritu Santo. Zoosystema 31, 453–469.

Ozawa, T., Köhler, F., Reid, D.G., Glaubrecht, M., 2009. Tethyan relicts on continental coastlines of the northwestern Pacific Ocean and Australasia: molecular phylogeny and fossil record of batillariid gastropods (Caenogastropoda, Cerithioidea). Zool Scr 38, 503–525.

Pagel, M., Meade, A., 2006. Bayesian analysis of correlated evolution of discrete characters by reversible-jump Markov chain Monte Carlo. Am Nat 167, 808–825.

Palmer, M.A., 1988. Dispersal of marine meiofauna - a review and conceptual model explaining passive tansport and active recruitment. Marine Ecology-Progress Series 48, 81–91.

Paradis, E., Claude, J., Strimmer, K., 2004. APE: analyses of phylogenetics and evolution in R language. Bioinformatics 20, 289–290.

Ponder, W.F., 1986. Glacidorbidea (Glacidorbacea, Basommatophora), a new family and superfamily of operculate fresh-water gastropods. Zool J Linn Soc 87, 53–83.

Posada, D., 2008. jModelTest: Phylogenetic model averaging. Mol Biol Evol 25, 1253–1256.

Rankin, J.J., 1979. A freshwater shell-less mollusc from the Caribbean: structure, biotics and contribution to a new understanding of the Acochlidioidea. R Ont Mus Life Sci Contrib 116, 1–123.

Ree, R.H., Moore, B.R., Webb, C.O., Donoghue, M.J., 2005. A likelihood framework for inferring the evolution of geographic range on phylogenetic trees. Evolution 59, 2299–2311.

Ree, R.H., Smith, S.A., 2008. Maximum likelihood inference of geographic range evolution by dispersal, local extinction, and cladogenesis. Syst Biol 57, 4–14.

Reid, D.G., Dyal, P., Williams, S.T., 2010. Global diversification of mangrove fauna: a molecular phylogeny of *Littoraria* (Gastropoda: Littorinidae). Mol Phylogenet Evol 55, 185–201.

Remane, A., 1940. Einführung in die zoologische Ökologie der Nord-und Ostsee. In: Grimpe, G., Wagler, E. (Eds.), Die Tierwelt der Nord-und Ostsee. Geest & Portig, Leipzig, pp. 1–238.

Rundell, R.J., Leander, B.S., 2010. Masters of miniaturization: Convergent evolution among interstitial eukaryotes. Bioessays 32, 430–437.

Salvini-Plawen, L.v., Steiner, G., 1996. Synapomorphies and plesiomorphies in higher classification of Mollusca. In: Taylor, J. (Ed.), Origin and evolutionary radiation of the Mollusca. Oxford University Press, Oxford, pp. 29–51.

Schrödl, M., 2006. Techniques for collecting interstitial opisthobranchs. In: Museum, S.S.F.A. (Ed.), Sydney, p. http://www.seaslugforum.net/factsheet/inteextr

Schrödl, M., Jörger, K.M., Klussmann-Kolb, A., Wilson, N.G., 2011a. Bye bye ‘Opisthobranchia’! A review on the contribution of mesopsammic sea slugs to euthyneuran systematics. Thalassas 27, 101–112.

Schrödl, M., Jörger, K.M., Wilson, N.G., 2011b. A reply to Medina et al. (2011): Crawling through time: Transition of snails to slugs dating back to the Paleozoic based on mitochondrial phylogenomics. Marine Genomics 4, 301–303.

Schrödl, M., Neusser, T.P., 2010. Towards a phylogeny and evolution of Acochlidia (Mollusca: Gastropoda: Opisthobranchia). Zool J Linn Soc 158, 124–154.

Smith, B.J., Stanisic, J., 1998. Pulmonata. Introduction. In: Beesley, P.L., Ross, G.J.B., Wells, A. (Eds.), Mollusca: the southern synthesis. Fauna of Australia. CSIRO Publishing, Melbourne, pp. 1037–1061.

Solem, A., 1985. Origin and diversification of pulmonate land snails. In: Trueman, E.R., Clarke, M.R. (Eds.), The Mollusca. Evolution. Academic Press, New York, pp. 269–293.

Solem, A., Yochelson, E.L., 1979. North American Paleozoic land snails, with a summary of other Paleozoic nonmarine snails. Geol Surv Prof Pap 1072, 1–39.

Sommerfeldt, N., Schrödl, M., 2005. Microanatomy of *Hedylopsis ballantinei*, a new interstitial acochlidian gastropod from the Red Sea, and its significance for phylogeny. J Molluscan Stud 71, 153–165.

Spalding, M.D., Fox, H.E., Halpern, B.S., McManus, M.A., Molnar, J., Allen, G.R., Davidson, N., Jorge,Z.A., Lombana, A.L., Lourie, S.A., Martin, K.D., McManus, E., Recchia, C.A., Robertson, J., 2007. Marine ecoregions of the world: A bioregionalization of coastal and shelf areas. Bioscience 57, 573-583.

Stadler, T., 2011a. Mammalian phylogeny reveals recent diversification rate shifts. Proc Natl Acad Sci USA 108, 6187–6192.

Stadler, T., 2011b. TreePar in R - Estimating diversification rates in phylogenies. Available at http://cran.r-project.org/web/packages/TreePar/index.html.

Stöger, I., Schrödl, M., 2013. Mitogenomics does not resolve deep molluscan relationships (yet?). Mol Phylogenet Evol 69, 376–392.

Stöger, I., Sigwart, J.D., Kano, Y., Knebelsberger, T., Marshall, B.A., Schwabe, E., Schrödl, M., 2013. The continuing debate on deep molluscan phylogeny: Evidence for Serialia (Mollusca, Monoplacophora plus Polyplacophora). Hindawi Biomed Res Int. doi.org/10.1155/2013/407072.

Swedmark, B., 1964. The interstitial fauna of marine sand. Biological Reviews 39, 1–42.

Swedmark, B., 1968. The biology of interstitial Mollusca. Symp Zool Soc London 22, 135–149.

Swennen, C.K., Buatip, S., 2009. *Aiteng ater*, new genus, new species, an amphibous and insectivorous sea slug that is difficult to classify (Mollusca: Gastropoda: Opisthobranchia: Sacoglossa(?): Aitengidae, new family). Raffles Bull Zool 57, 495–500.

Talavera, G., Castresana, J., 2007. Improvement of phylogenies after removing divergent and ambiguously aligned blocks from protein sequence alignments. Syst Biol 56, 564–577.

Tardy, J., 1970. Contribution à l’étude des métamorphoses chez les nudibranches. Ann Sci Nat Zool Biol Anim 12, 299–370.

Tracey, S., Todd, J.A., Erwin, D.H., 1993. Mollusca: Gastropoda. In: Benton, M.J. (Ed.), The fossil record. Chapman and Hall, London, pp. 131–167.

Vermeij, G.J., 1991. Anatomy of an invasion - the trans-arctic interchange. Paleobiology 17, 281–307.

Vermeij, G.J., 2005. From Europe to America: Pliocene to recent trans-Atlantic expansion of cold-water North Atlantic molluscs. Proc. R. Soc. B-Biol. Sci. 272, 2545–2550.

Vermeij, G.J., Dudley, R., 2000. Why are there so few evolutionary transitions between aquatic and terrestrial ecosystems? Biol J Linn Soc 70, 541–554.

Vermeij, G.J., Wesselingh, F.P., 2002. Neogastropod molluscs from the Miocene of western Amazonia, with comments on marine to freshwater transitions in molluscs. J Paleontol 76, 265–270.

Wägele, H., Klussmann-Kolb, A., 2005. Opisthobranchia (Mollusca, Gastropoda) - more than just slimy slugs. Shell reduction and its implications on defence and foraging. Front Zool 2, 1–18.

Wägele, H., Klussmann-Kolb, A., Verbeek, E., Schrödl, M., 2014. Flashback and foreshadowing—a review of the taxon Opisthobranchia. Org Divers Evol 14, 133–149.

Wägele, H., Klussmann-Kolb, A., Vonnemann, V., Medina, M., 2008. Heterobranchia I: The Opisthobranchia. In: Ponder, W.F., Lindberg, D. (Eds.), Phylogeny and Evolution of the Mollusca. University of California Press, Berkeley, pp. 385–408.

Wawra, E., 1974. The rediscovery of *Strubellia paradoxa* (Strubell) (Gastropoda: Euthyneura: Acochlidiacea) on the Solomon Islands. The Veliger 17, 8–10.

Wawra, E., 1979. *Acochlidium sutteri* nov. spec. (Gastropoda, Opisthobranchia, Acochlidiacea) von Sumba, Indonesien. Ann Nathist Mus Wien Ser B Bot Zool 82 595–604.

Wawra, E., 1987. Zur Anatomie einiger Acochlidia (Gastropoda, Opisthobranchia) mit einer vorläufigen Revision des Systems und einem Anhang über Platyhedylidae (Opisthobranchia, Ascoglossa). Universität Wien, Wien.

Westheide, W., 1982. *Microphthalmus hamosus* sp. n. (Polychaeta, Hesionidae) - an example of evolution leading from the interstital fauna to a macrofaunal interspecific relationship. Zool Scr 11, 189–193.

Westheide, W., 1987. Progenesis as a principle in meiofauna evolution. J Nat Hist 21, 843–854.

White, T.R., Conrad, M.M., Tseng, R., Balayan, S., Golding, R., de Frias Martins, A.M., Dayrat, B.A., 2011. Ten new complete mitochondrial genomes of pulmonates (Mollusca: Gastropoda) and their impact on phylogenetic relationships. BMC Evol Biol 11.

Williams, S.T., 2007. Origins and diversification of Indo-West Pacific marine fauna: evolutionary history and biogeography of turban shells (Gastropoda, Turbinidae). Biol J Linn Soc 92, 573–592.

Williams, S.T., Duda, T.F., Jr., 2008. Did tectonic activity stimulate Oligo-Miocene speciation in the Indo-West Pacific? Evolution 62, 1618–1634.

Worsaae, K., Kristensen, R.M., 2005. Evolution of interstitial Polychaeta (Annelida). Hydrobiologia 535, 319–340.

Worsaae, K., Rouse, G.W., Kristensen, R.M., Edgecombe, G.D., Sterrer, W., Pleijel, F., Giribet, G., 2008. Origin of interstitial Annelida. J Morphol 269, 1462-1462.

Worsaae, K., Sterrer, W., Kaul-Strehlow, S., Hay-Schmidt, A., Giribet, G., 2012. An anatomical description of a miniaturized acorn worm (Hemichordata, Enteropneusta) with asexual reproduction by paratomy. PLoS ONE 7.

Yu, Y., A.J., H., X.J., H., 2012. RASP (Reconstruct Ancestral State in Phylogenies) 2.1b. http://mnh.scu.edu.cn/soft/blog/RASP.

Zapata, F., Wilson, N.G., Howison, M., Andrade, S.C.S., Jörger, K.M., Schrödl, M., Goetz, F.E., Giribet, G., Dunn, C.W., 2014. Phylogenomic analyses of deep gastropod relationships reject Orthogastropoda. Proceedings of the Royal Society B: Biological Sciences 281.

